# The modifiers that cause changes in gene essentiality

**DOI:** 10.1101/2025.03.06.641712

**Authors:** Amandine Batté, Núria Bosch-Guiteras, Carles Pons, Marina Ota, Maykel Lopes, Sushma Sharma, Nicolò Tellini, Claire Paltenghi, Michelle Conti, Kwan Ting Kan, Uyen Linh Ho, Michaël Wiederkehr, Jonas Barraud, Mark Ashe, Patrick Aloy, Gianni Liti, Andrei Chabes, Leopold Parts, Jolanda van Leeuwen

## Abstract

The phenotype of a mutation often differs across genetically distinct individuals. In the most extreme case, a gene can be essential for viability in one genetic background, but dispensable in another. Although genetic context-dependency of mutant phenotypes is frequently observed, the underlying causes often remain elusive. Here, we investigated the genetic changes responsible for differences in gene essentiality across 18 genetically diverse natural yeast strains. First, we identified 39 genes that were essential in the laboratory reference strain but not required for viability in at least one other genetic background, suggesting that the natural strain contained suppressor variants that could bypass the need for the essential gene. We then mapped and validated the causal bypass suppressor variants using bulk segregant analysis and allele replacements. Bypass suppression was generally driven by a single modifier gene that tended to differ between genetic backgrounds. The suppressors often indirectly counteracted the effect of deleting the essential gene, for instance by changing the transcriptome of a cell. Context-dependent essential genes and their bypass suppressors were frequently co-mutated across 1,011 yeast isolates and identified naturally occurring evolutionary trajectories. Overall, our results highlight the relatively high frequency of bypass suppression in natural populations, as well as the underlying variants and mechanisms. A thorough understanding of the causes of genetic background effects is crucial for the interpretation of genotype-to-phenotype relationships, including those associated with human disease.

## INTRODUCTION

The consequence of a mutation often depends on the genetic background in which it occurs. In extreme cases, a mutation can be lethal in one individual, while having no apparent effect in another. Such incomplete penetrance complicates the prediction of phenotype from genotype, thereby impeding disease diagnosis before the onset of symptoms (Riordan and Nadeau 2017). Incomplete penetrance can arise due to additional variants in the genetic background that exacerbate or reduce the impact of a given mutation (Genin et al. 2008, Harper et al. 2015). Identification of such modifying variants would increase our understanding of how this phenotypic variation arises and improve our ability to predict disease risk, yet, finding these variants in populations of genetically diverse patients is not trivial. As a result, modifiers have mainly been described for relatively common Mendelian disorders, such as cystic fibrosis and sickle cell disease (Steinberg and Sebastiani 2012, Ünlü et al. 2023).

The budding yeast *Saccharomyces cerevisiae* is a powerful model system for studying modifier mutations, due to its highly annotated genome and tractable genetics (Botstein and Fink 2011). Gene essentiality is arguably one of its most straight-forward phenotypes to analyze. The set of essential yeast genes was defined by individually deleting one copy of each gene in a diploid cell and testing for viability of haploid offspring carrying the deletion allele (Giaever et al. 2002). In total, 17% (1,105) of the ∼6,000 yeast genes were found to be essential for viability under nutrient-rich growth conditions in the laboratory strain S288C. Although essential genes tend to play highly conserved roles in a cell (Giaever et al. 2002, Rancati et al. 2018), genetic modifiers sometimes lead to a rewiring of cellular processes that bypasses the requirement for specific essential genes (Bosch-Guiteras and Van Leeuwen 2022). Comparison of two closely related budding yeast strains revealed that ∼6% of essential genes are uniquely essential in one strain background (Dowell et al. 2010). When comparing the essential gene set of *S. cerevisiae* to other yeast species, the percentage of differential essential genes increases from ∼12% for the closely related species *Saccharomyces uvarum* to ∼17% for the highly diverged *Schizosaccharomyces pombe* (Kim et al. 2010, Sanchez et al. 2019). Differences in gene essentiality are also common in more complex cells, as ∼25% of essential genes in any given human cell line could be classified as context-specific essential (Hart et al. 2015).

Although differences in gene essentiality or mutant fitness across yeast isolates are well documented (Dowell et al. 2010, Hou et al. 2019, Johnson et al. 2019, Caudal et al. 2022, Chen et al. 2022, Wang et al. 2022, Hale et al. 2024), the genetic variants responsible for the observed differences remain largely unknown. Some insights on the causes of context-dependent gene essentiality come from systematic studies that isolated spontaneous mutations that can bypass the requirement for an essential gene in a laboratory yeast strain (Liu et al. 2015, Chen et al. 2016, Van Leeuwen et al. 2020). The bypass suppressors often involved gain-of-function mutations in essential genes or in genes with a functional connection to the bypassed essential gene, such as shared complex or pathway membership (Van Leeuwen et al. 2020). Most essential genes could be suppressed by mutations in only one or two genes, suggesting that there are only a few fundamental ways of rewiring biological processes such that deletion of an essential gene can be tolerated (Van Leeuwen et al. 2020). However, the relevance of these findings for natural populations remains unclear, as long-term evolution may have resulted in more complex mechanisms of suppression that either involve mutations in multiple suppressor genes simultaneously or are caused by specific variants that are not readily obtained in the relatively small populations and short time scales used in a laboratory setting. Here, we explored the natural causes of variation in gene essentiality across 18 diverse *S. cerevisiae* strains, to determine the general principles of bypass suppression by natural variation.

## RESULTS

### Changes in gene essentiality are common among yeast isolates

To uncover differences in gene essentiality across diverse genetic backgrounds, we focused on genes that were essential in the laboratory strain S288C and searched for genetic contexts in which these genes were not required for viability. We used the Synthetic Genetic Array (SGA) approach to cross a collection of “query” mutant strains in the S288C background to 18 yeast isolates from various sources and geographic locations (“wild strains”; **Fig. 1A-B**), as well as an S288C control. The query strain collection consisted of haploid strains, each deleted for an essential gene in their genome but viable because of the presence of the essential gene on a counter-selectable plasmid (Van Leeuwen et al. 2020). The collection contained 2,539 query strains for 763 different essential genes (∼70% of all essential yeast genes), with 348 of these genes represented by multiple strains. The selected wild strains carried ∼30,000-80,000 variants compared to the S288C reference, and had, on average, two nonsynonymous and three silent mutations per gene (**Data S1**). FY4 was included as a negative control, as this strain was derived from S288C (Winston et al. 1995, Brachmann et al. 1998) and had only 244 genomic variants compared to the reference sequence (**Data S1**).

**Figure 1.**
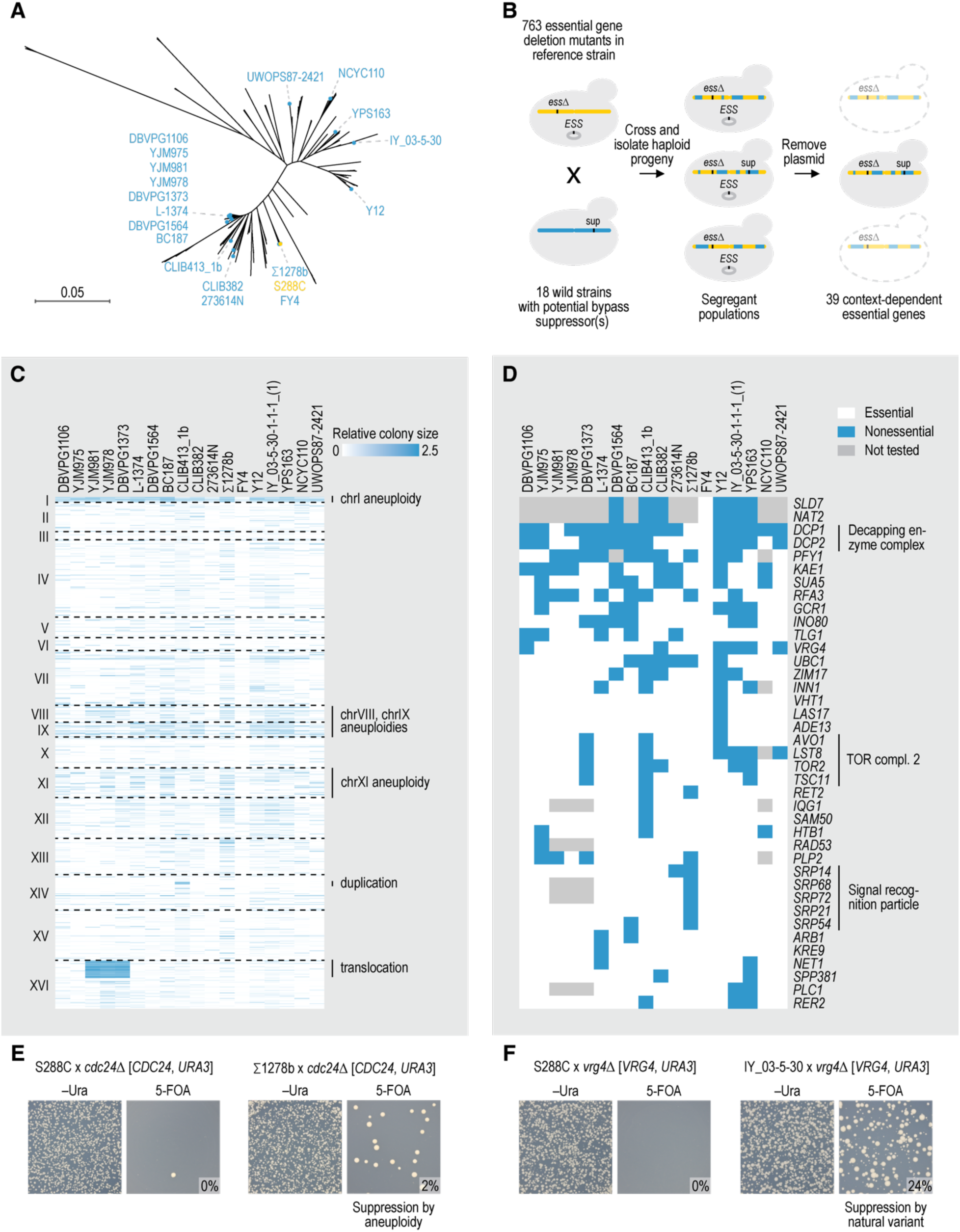
High-throughput identification of genetic context-dependent essential genes. (**A**) Phylogenetic tree of *Saccharomyces cerevisiae*, indicating the various wild strains (blue) and the S288C reference strain (yellow) that were used in this study. (**B**) Strategy for identifying genetic context-dependent essential genes. A collection of haploid strains in the reference background, each deleted for an essential gene in their genome but carrying it on a counter-selectable plasmid, was crossed to 18 wild strains, as well as an S288C control, and haploid segregant populations carrying both the deletion allele and the complementing plasmid were isolated. Growth in the absence of the essential gene was subsequently assessed by counter-selecting against the plasmid. (**C**) Relative colony size of haploid segregant populations lacking an essential gene. Each row represents a different essential gene, sorted by genomic coordinate. Columns correspond to the wild yeast strains used in the crosses and are ordered by genetic similarity. Large genomic determinants that affected gene essentiality measurements are indicated. (**D**) Identified context-dependent essential genes and the wild strain crosses in which the genes were nonessential are shown. Wild strains are sorted by genetic similarity. (**E-F**) Examples of the secondary validation assay in case of suppression of the essential gene deletion mutant by either a spontaneous aneuploidy that provides an extra copy of the essential gene (E) or by natural variants (F). *cdc24*Δ [*CDC24*, *URA3*] and *vrg4*Δ [*VRG4*, *URA3*] strains in the reference background were crossed to both S288C and a wild strain. Growth of haploid single colony progeny was determined in the presence (–Ura) and absence (5-FOA) of the plasmid carrying the essential gene. The percentage of colonies that were obtained on 5-FOA media compared to –Ura media is indicated. IY_03-5-30 = IY_03-5-30-1-1-1_(1).

Each query-wild strain cross was driven through meiosis, and pools of haploid segregant progeny carrying both the essential gene deletion allele and the plasmid with the essential query gene were selected. Cells were subsequently transferred to a medium that selected against the plasmid, to assess for growth in the absence of the essential gene (**Fig. 1B**, **Data S2**). To be able to confidently distinguish pools that contained viable segregants from those that did not, we included four technical replicates of each cross and a mean of six biological replicates for each essential gene. The technical replicates allowed removal of false positive cases of cell growth that can arise due to spontaneous mutations in the counter-selectable marker that prevent removal of the complementing plasmid. We considered a deletion mutant suppressed in a genetic background when, after filtering false positive cases, > 50% of independently generated wild segregant pools showed growth, and the corresponding S288C control progeny did not (Methods).

Our screen identified large structural variants that could provide an extra copy of the essential query genes located in the affected regions, thereby masking the lethality of the essential gene deletion mutant (**Fig. 1C**). The strains DBVPG1373, YJM978, and YJM981 harbor a translocation between chromosomes VIII and XVI (Perez-Ortin et al. 2002). When these strains are crossed to the reference strain, 25% of the progeny carries a duplication of a substantial part of the left arm of chromosome XVI (Parts et al. 2021). As expected, all segregant pools of strains carrying the translocation crossed to essential gene deletion mutants located in the duplicated region showed growth in our screen, due to the presence of a complementing copy of the essential gene in 25% of the segregants (**Fig. 1C**, **S1A**). Similarly, four adjacent genes on the left arm of chromosome XIV appeared nonessential in crosses with CLIB413_1b, due to a duplication of the region between *YNL251C* and *YNL243W* in this background (**Fig. 1C**, **S1B**). Finally, most segregant pools of essential genes located on chromosome I showed growth in our screen. A similar, though less extreme, trend was observed for chromosomes VIII, IX, and XI (**Fig. 1C**, **S1C**). Aneuploidies of chromosome I, VIII, IX, and XI are frequently observed in natural yeast isolates, suggesting that they may occur spontaneously at a relatively high rate and are generally well tolerated (Peter et al. 2018, Scopel et al. 2021). Although the wild yeast strains that were used in the crosses did not carry aneuploidies of chromosomes I, VIII, IX, or XI, we suspected that these aneuploidies could be selected for during our screen, masking the lethality of essential gene deletion alleles located on these chromosomes.

To distinguish between growth in segregant pools due to the deleted gene being nonessential in the presence of natural variants and growth due to the presence of a second copy of the essential gene provided by an aneuploidy, we examined the viability of hundreds of single colony progeny from each segregant pool in which growth was observed. We compared the number of obtained colonies per pool both in the presence and absence of the plasmid covering the essential gene deletion allele and calculated the percentage of progeny that was viable without the plasmid (**Fig. 1E-F**, **Data S3**). Spontaneous aneuploidies are expected to be relatively rare and thus only a small fraction of progeny will be viable without the complementing plasmid in such cases (**Fig. 1E**). By contrast, bypass suppression by standing variation would result in 12.5% to 50% viable progeny in the absence of the essential gene plasmid for cases involving between one and three modifier loci (**Fig. 1F**). We therefore set a stringent threshold for calling a gene context-dependent essential by requiring that at least 10% of segregant progeny for a given cross were viable on media selecting against the complementing plasmid (Discussion) and verified absence of the essential gene in the viable progeny by sequencing (**Data S3**). Overall, 11% of the potential conditional essential genes that were located on one of the frequently aneuploid chromosomes validated as context-dependent essential, as did 55% of the genes located on one of the other chromosomes (**Data S3**). As expected, all genes that were essential in S288C were also required for viability in the negative control strain FY4, which was derived from S288C. In total, we identified 39 context-dependent essential genes that were nonessential in the presence of natural variants from at least one wild strain (5% of tested essential genes, **Fig. 1D**, **Data S4**).

### Consistency in context-dependent gene essentiality across studies

A previous study individually deleted ∼5100 genes in the Σ1278b strain and uncovered 13 genes that were essential in S288C but not in Σ1278b (Dowell et al. 2010). Eight of these genes were included in our screen. Reassuringly, we identified six of the eight genes as nonessential in the Σ1278b cross (*PFY1*, *PLP2*, *SRP14*, *SRP21*, *SRP68*, and *SRP72*; 262-fold enrichment over expectation, *p* < 0.0005, Fisher’s exact test). When tested individually, we did obtain viable progeny for deletion mutants of the remaining two genes in a Σ1278b cross, but they were not identified as nonessential in our screen due to either a severe fitness defect of the deletion mutant (*UBC1*) (Dowell et al. 2010) or high variation in colony size among technical replicates (*RET2*; **Data S2**). Furthermore, we found two additional genes (*SRP54* and *RFA3*) that were not required for viability in the Σ1278b cross, but that had not been previously described as nonessential in this background. We conclude that our approach for discovering context-dependent essential genes results in a high level of experimental accuracy that is comparable to individually deleting genes in a wild background.

Five of the 39 identified context-dependent essential genes also showed variation in viability in a study examining gene essentiality across 15 natural yeast isolates that were not included in our screen (**Fig. S2A**; 9-fold enrichment over expectation, *p* < 0.005, Fisher’s exact test) (Chen et al. 2022). Moreover, consistent with their context-dependency in our experiments, essential genes that were nonessential in at least one of the tested wild strains frequently carried homozygous loss-of-function mutations in a collection of 1,011 natural yeast isolates (**Fig. 2A**) (Peter et al. 2018). We further compared our identified context-dependent essential genes with those previously found to be bypassed by spontaneous mutations in the S288C reference strain, and found significant overlap between both gene sets (**Fig. 2B**, **S2A**) and their properties (**Fig. S2B**) (Van Leeuwen et al. 2020, Pons and Van Leeuwen 2023). In total, 37 out of 39 identified context-dependent essential genes (95%) showed evidence for dispensability in at least one other study (**Data S5**; 2-fold enrichment over expectation, *p* < 0.0005, Fisher’s exact test). Based on these results, we updated a previously defined list of core essential yeast genes (Van Leeuwen et al. 2020). Of the genes that were classified as essential in S288C, 436 genes are invariably required for cell viability in *S. cerevisiae*, 264 genes vary in essentiality across genetic backgrounds, and 357 genes remain to be classified (**Data S5**; Methods).

**Figure 2.**
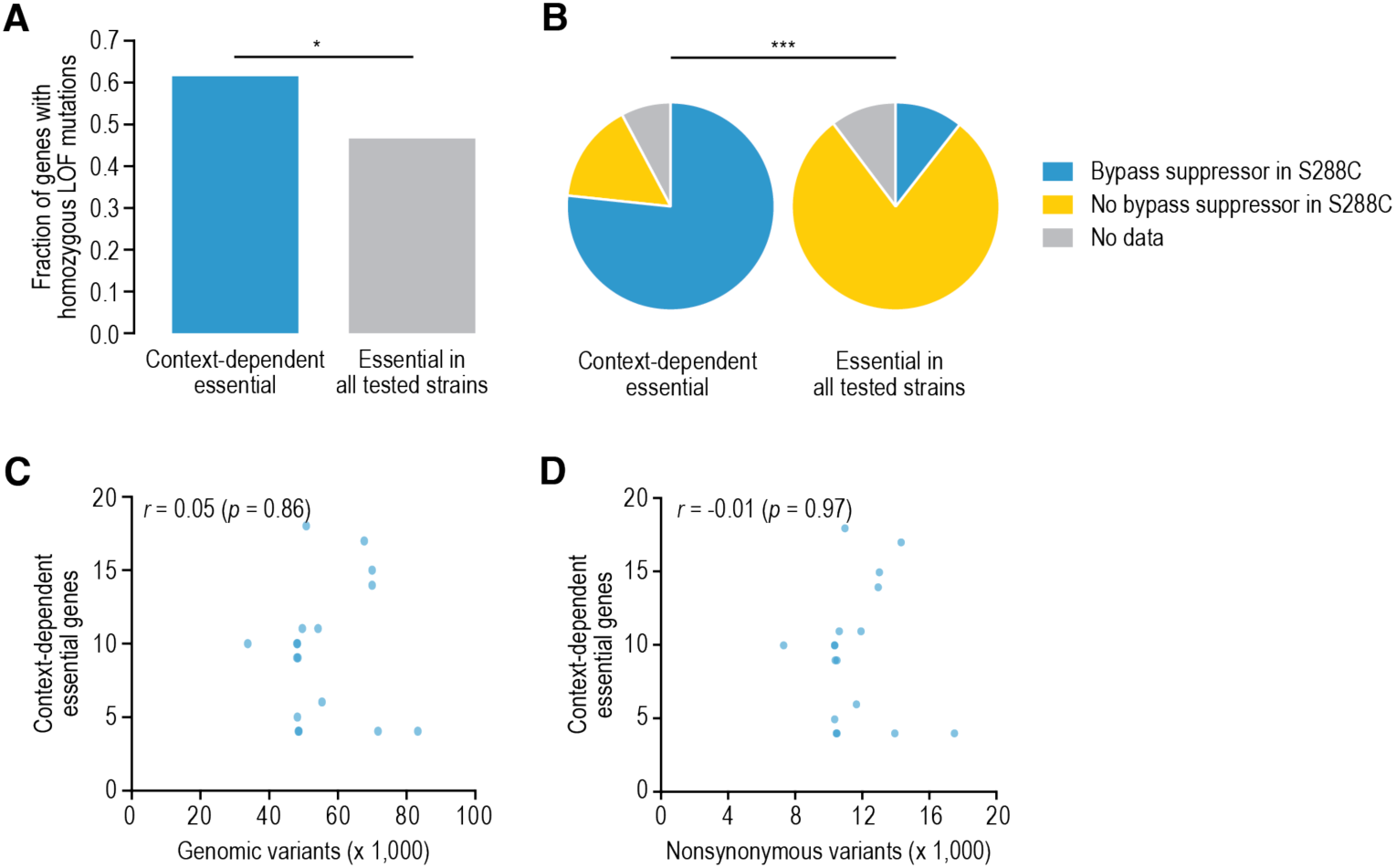
Properties of context-dependent essential genes. (**A**) Fraction of context-dependent essential genes (*N* = 39) or genes that were essential in all tested wild strains (*N* = 724) that carried homozygous loss-of-function (LOF) mutations in one or more of 1,011 natural yeast isolates (Peter et al. 2018). Strains that were used in our screen were excluded from the analysis. * *p* < 0.05, one-sided Fisher’s exact test. (**B**) Overlap of genetic-context dependent essential genes discovered in this study with either essential genes for which spontaneous bypass suppressor mutations have been found in S288C or essential genes for which no bypass suppressors could be isolated in the reference strain (Van Leeuwen et al. 2020). *** *p* < 0.0005, **χ**^2^ test. (**C**-**D**) The number of context-dependent essential genes identified in a given wild strain is plotted against the number of variants in the wild genome compared to the S288C reference, considering either all variants (C) or nonsynonymous variants only (D). Pearson’s correlation coefficients and the corresponding *p* values are indicated.

### Limited correlation between genetic relatedness and changes in gene essentiality

To gain global insights into the genetic underpinnings of context-dependent gene essentiality, we first explored patterns of the phenotype across genetic backgrounds. Excluding the negative control strain FY4, the number of context-dependent essential genes ranged from 4 to 18 per genetic background, with an average of nine genes that were essential in S288C but not in a given background. There was no significant correlation between the number of detected context-dependent essential genes in a genetic background and the number of variants compared to the reference (**Fig. 2C-D**).

Consistent with their overlap with other studies using different wild strains (**Fig. S2A**), most identified context-dependent essential genes (29 of 39 genes, 74%) were nonessential in multiple genetic backgrounds in our study, with six genes being classified as dispensable in over half of the tested backgrounds (**Fig. 1D**). These six genes span a range of functions and include the decapping enzyme complex genes *DCP1* and *DCP2* and *PFY1* encoding for profilin. Somewhat surprisingly, the wild strains that shared context-dependent essential genes were generally not closely related (**Fig. 1D**), and context-dependent essentiality was only modestly consistent with genetic relatedness (**Fig. S2C**), suggesting that context-dependent essential genes can often be bypassed by multiple independent suppressor variants.

### Genetic architecture of bypass suppression

To identify the natural variants that were responsible for the change in gene essentiality, we performed bulk segregant analysis (Liti and Louis 2012) on query strains of 37 out of 39 context-dependent essential genes crossed to a genetic background in which the gene was nonessential. For seven context-dependent essential genes, we included two different genetic backgrounds that ideally came from two different clades, for a total of 44 crosses. We isolated meiotic progeny both in the presence and absence of the plasmid covering the essential gene deletion allele, sequenced the populations, and compared variant allele frequencies between populations with and without the plasmid (**Data S6**). Most segregant pools (35 of 44 pools, 80%) showed regions of specific selection for wild sequence in the absence of the plasmid (**Fig. 3A-B**, **Data S6**, **S7**), with an average of two suppressor loci per cross. In the remaining nine pools, bypass suppression may be caused by variants in regions that were excluded from the analysis, such as the mitochondrial genome and regions around selection markers (Methods). Essential genes that encoded members of the same protein complex showed similar suppressor loci when crossed to the same wild strain (**Fig. 3B**; *SRP21*, *SPR54*, *SPR68*, and *SPR72*; or *AVO1*, *LST8*, *TOR2*, and *TSC11*). By contrast, when the same essential gene deletion mutant was crossed to different wild strains, suppressor loci were often different (e.g., *HTB1*, *PFY1*), consistent with the observed patterns of context-dependent gene essentiality that suggested that bypass suppressor variants can differ between genetic backgrounds (**Fig. 1D**, **S2C**).

**Figure 3.**
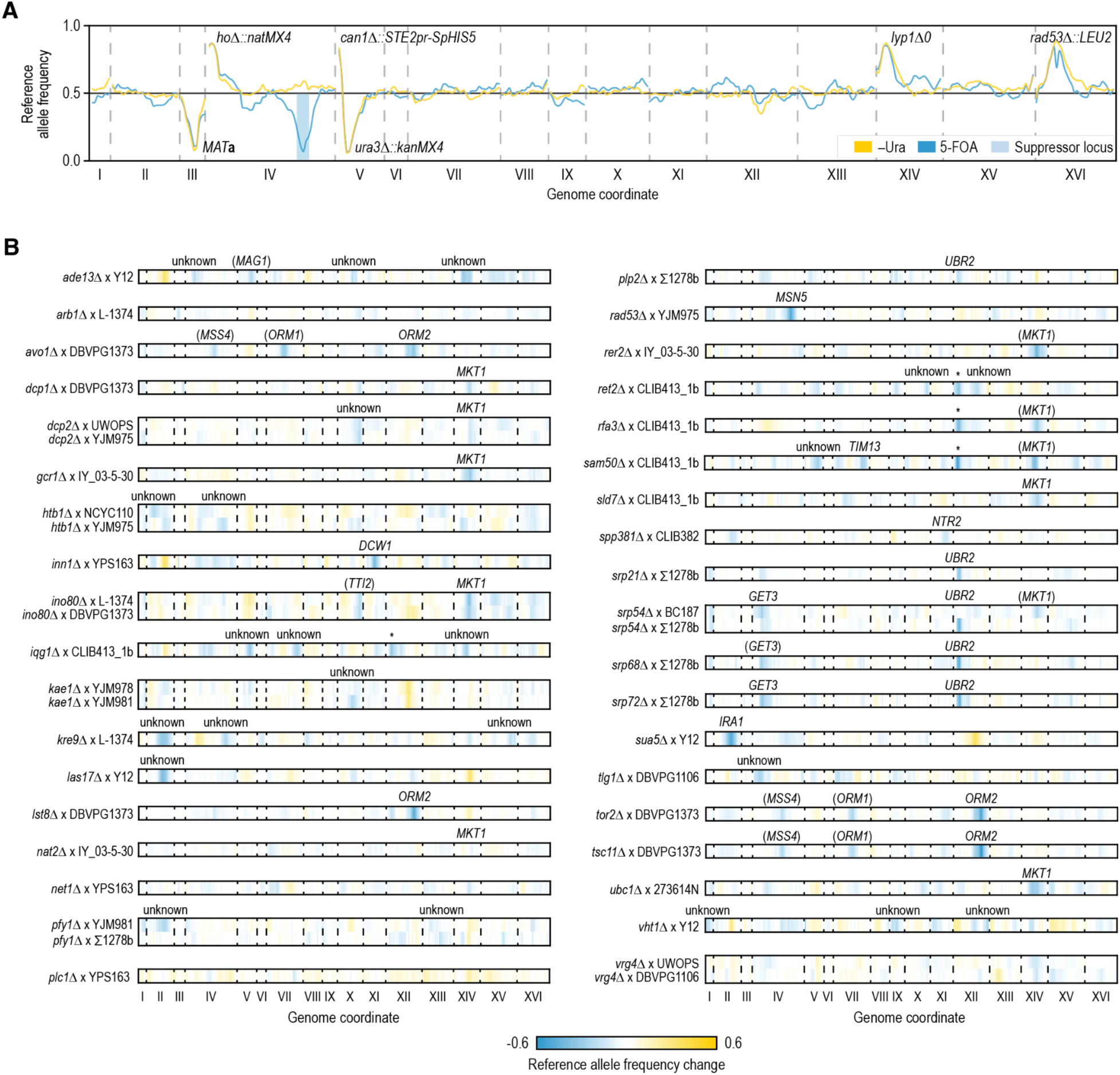
Mapping suppressor loci by sequencing segregant pools. (**A**) Example of the mapping results. Reference allele frequency along the yeast genome in progeny of a cross between a *rad53*Δ [*RAD53*, *URA3*] S288C strain and YJM975, either in the presence of the plasmid carrying *RAD53* (–Ura, yellow) or in the absence of the plasmid (5-FOA, cyan). Loci that carried selection markers that were used in the experiment are labeled. The difference in allele frequency between the two conditions is used in the mapping. (**B**) Mapping results for segregant pools involving the indicated essential gene deletion mutants and wild strains. The change in S288C allele frequency between the meiotic progeny isolated in the presence and in the absence of the plasmid covering the essential gene deletion allele is plotted by genomic coordinate. Causal bypass suppressor genes are indicated for regions that show selection for wild strain sequence. Genes in brackets have not been validated experimentally as bypass suppressors. IY_03-5-30 = IY_03-5-30-1-1-1_(1); UWOPS = UWOPS87-2421; * = *PPR1* locus that shows selection for S288C sequence in –Ura samples of the indicated crosses.

Progeny of mutant strains crossed with CLIB413_1b showed strong selection for S288C sequence on the left arm of chromosome XII, around the gene *PPR1*, when grown on media lacking uracil (**Fig. 3B**, **S3A**). This media was used to select for the *URA3* marker on the plasmid carrying the essential gene in control samples. Ppr1 positively regulates transcription of *URA3*, which encodes an enzyme that converts orotidine-5’-phosphate into uridine monophosphate, a crucial step in uracil biosynthesis (Flynn and Reece 1999). CLIB413_1b carries a G95A missense mutation in *PPR1* that is predicted to be deleterious (Vaser et al. 2016), suggesting that the S288C allele of *PPR1* is beneficial when uracil is not provided in the media. Reassuringly, selection against the plasmid by 5-fluoroorotic acid (5-FOA), which is converted by Ura3 into the toxic 5-fluorouridine (Boeke et al. 1984), still worked efficiently in these strains, as no reads mapping to the plasmid were detected in the sequencing data of these samples.

In 14 out of 44 crosses, sequence from the S288C reference background was selected in the absence of the complementing plasmid, often in addition to loci with strong selection for wild sequence (**Fig. 3B**, **Data S7**). This could suggest that a combination of both wild and S288C alleles is needed to bypass the essential gene. To test this, we selected 14 genes that showed different selection patterns in the bulk segregant analysis, i.e. either selection for wild sequence only, for S288C sequence only, for both, or for neither. We directly knocked out these genes in the wild strain, as well as an S288C control, and tested viability. For 12 out of 14 tested genes (86%), many viable colonies were obtained in the wild background, but not in S288C, confirming that essential gene bypass is driven by variants present in the wild genome (**Fig. S3B**). In crosses that show selection for S288C sequence, the S288C alleles may further improve the fitness of the bypassed strains. For the remaining two genes (*NET1* and *RER2*), < 20% of the colonies lacking the gene survived in the wild background. Neither of these genes, however, showed significant selection for S288C sequence in the bulk segregant analysis. Possibly, nonchromosomal elements contribute to the bypass of these genes (Edwards et al. 2014). Overall, we conclude that genomic variants present in the wild background are sufficient to bypass the essential gene in the vast majority of cases.

### Bypass suppressor genes tend to act in isolation

Each of the mapped suppressor loci harbored multiple genes and variants. To identify the genes that caused bypass of the essential gene, we replaced candidate suppressor genes in the reference strain background with the corresponding wild sequence and tested for viability of the resulting strains in the absence of the essential gene. We selected candidate suppressor genes within each locus based on known functional connections (Engel et al. 2024) or physical or genetic interactions (Oughtred et al. 2021) with the bypassed essential gene. We also included *MKT1* as a candidate suppressor, as it is known to modify many different phenotypes (Fay 2013, Albert et al. 2014, Parts et al. 2014, Albert et al. 2018). Furthermore, we included six candidate genes in loci with weaker selection (difference in reference allele frequency of 0.14 to 0.17) that did not meet our criteria for calling suppressor loci (difference in reference allele frequency > 0.20), one candidate that was strongly selected both in the presence and absence of the plasmid carrying the essential gene (*MKT1* in *nat2*Δ; **Fig. S3C**), and one candidate gene that was genetically linked to the essential gene deletion allele (*MKT1* in *dcp2*Δ). The wild allele of one out of six (17%) of the candidate genes in a locus with weaker selection and 21 out of 47 (45%) of the candidates in strong suppressor loci could bypass the requirement for the essential gene, validating our mapping strategy (**Fig. S4**). In total, including the *MKT1 nat2*Δ and *MKT1 dcp2*Δ pairs, we identified causal bypass suppressor genes for 24 loci, bypassing 21 essential genes (**Fig. 4A**, **Data S7**).

**Figure 4.**
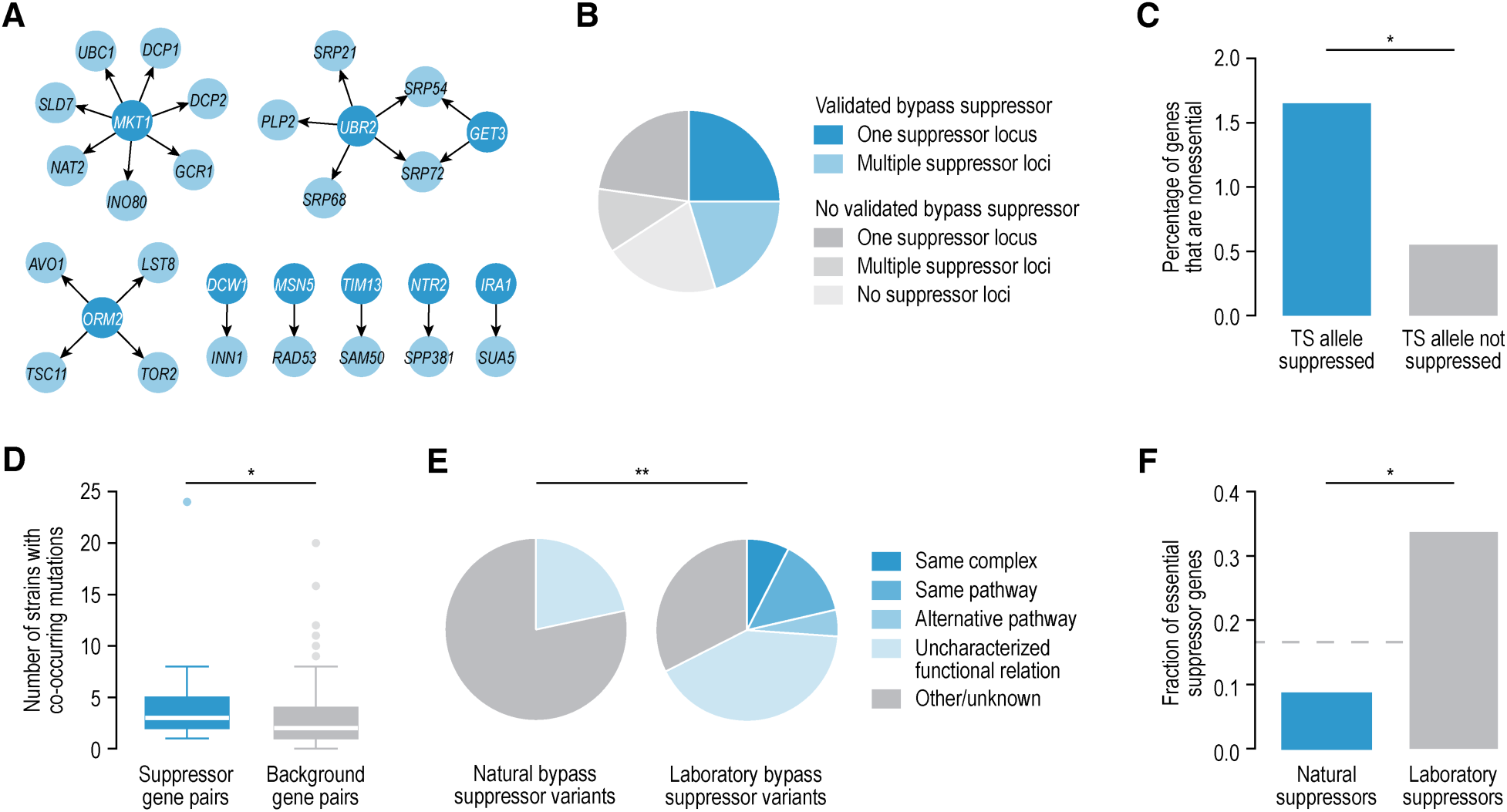
Characterization of natural bypass suppression variants. (**A**) Identified bypass suppression interactions. Interactions are represented as arrows that point from the bypass suppressor gene to the bypassed essential gene. (**B**) The fraction of crosses with mapping results (Fig. 3), for which we did or did not validate a bypass suppressor gene, divided by the number of mapped suppressor loci. (**C**) Percentage of essential S288C genes that are nonessential in a given background, for genes with TS alleles that either could (*N* = 431) or could not (*N* = 3,284) be suppressed in the same background. (**D**) Plotted are the number of yeast strains among 1,011 natural yeast isolates (Peter et al. 2018) in which a context-dependent essential gene carries homozygous loss-of-function mutations or deletions and the corresponding bypass suppressor gene is deleted, duplicated, or carries loss-of-function mutations (*N* = 23 pairs), compared to the co-occurrence of such mutations in randomized gene pairs (*N* = 166). Strains that were used in our screen were excluded from the analysis. (**E**) Distribution of suppression interactions across different mechanistic suppression classes, for spontaneous suppressor mutations of essential gene deletion mutants identified in the reference strain S288C in a laboratory setting (Van Leeuwen et al. 2020) (*N* = 130) or for natural variant suppressors identified in this study (*N* = 23). (**F**) The fraction of suppressor mutations that map to an essential gene, for the natural variant suppressors of essential gene deletion mutants identified in this study (*N* = 23) or for spontaneous suppressors of essential gene deletion mutants identified previously in the laboratory strain S288C (Van Leeuwen et al. 2020) (*N* = 130). Gray line, fraction of all yeast genes that is essential (0.17). Statistical significance was determined using Fisher’s exact (B,C,F), Mann-Whitney U (D), or **χ**^2^ (E) tests. * *p* < 0.05, ** *p* < 0.005, *** *p* < 0.0005.

The 24 validated bypass suppressors all acted in isolation, despite detecting multiple loci with selection for wild sequence in 10 of the 24 crosses with a validated suppressor (**Fig. 3B**, **4B**, **Data S7**). In the cases where multiple suppressor loci were mapped, but one was sufficient for bypass suppression to occur, the additional loci may contribute to the overall fitness of the strain. Indeed, although an *MKT1-BC187* allele could not bypass *srp54*Δ in the S288C background, it could slightly improve the fitness of viable *srp54Δ GET3-BC187* strains (**Fig. S5A**). For an additional 15 crosses, none of the introduced wild alleles could suppress the essential gene deletion mutant in the S288C background (**Fig. 4B**, **S4**, **Data S7**). It is possible that in these cases, multiple loci need to simultaneously express wild alleles for bypass suppression to occur. However, when we combined wild alleles of suppressor candidates from multiple loci for *IQG1* and *VHT1*, the genes remained inviable in S288C (**Fig. S5B**). Furthermore, in 10 out of 15 cases only a single suppressor locus was mapped (**Fig. 4B**), suggesting that cases that involve multiple unlinked bypass suppressor genes are rare.

### Little overlap between suppressors of hypomorphic and deletion alleles

We previously identified natural variants that could improve the fitness of essential gene temperature sensitive (TS) alleles at elevated temperature (Parts et al. 2021). When comparing the essential S288C genes that were nonessential in a given wild strain to those with a TS allele that could be suppressed in the same background, we found significant overlap between both datasets (**Fig. 4C**). Nonetheless, only 1.7% of the genes that had a TS allele that could be suppressed, were also nonessential in that genetic background. Although TS allele suppressors thus generally cannot suppress a deletion allele of the same gene (Discussion), exceptions occur. One such exception is *TSC11*, for which we had mapped suppressor variants of two TS alleles (Parts et al. 2021) as well as the deletion allele (this study) in crosses with the wild strain DBVPG1373. Comparison of the suppressor loci showed that in both cases three loci were identified on chromosomes IV, VII, and XII. However, selection of the chromosome XII locus was substantially stronger in the *tsc11*Δ cross than in the TS allele crosses, while the other two suppressor loci showed comparable selection (**Fig. S6A**). We confirmed that the wild allele of *ORM2*, that is located within the chromosome XII locus, is sufficient to bypass *tsc11*Δ (**Fig. S4**), consistent with its strong selection in the *tsc11*Δ cross. Thus, even in rare cases where suppressors of TS and deletion alleles overlap, the relative contribution of a particular gene to the suppression phenotype can differ.

### Bypass suppressors may affect evolutionary trajectories in natural yeast populations

Next, we explored the impact of the identified bypass suppressor genes on natural evolution. The genomes of 1,011 budding yeast isolates (Peter et al. 2018) are strongly depleted for homozygous loss-of-function variants in genes that are essential in the laboratory strain S288C (**Fig. S6B**). When variants that can act as bypass suppressors are present in a genome, selection against loss-of-function mutations in the suppressible essential genes could be reduced. To test this, we asked whether context-dependent essential genes and their bypass suppressors were more often co-mutated in the 1,011 wild genomes than expected by chance (Pons and Van Leeuwen 2023). Specifically, we investigated whether in strains in which the function of an essential gene was impaired through loss-of-function mutations or gene deletion events, the identified bypass suppressor gene carried impactful mutations too. For many of the suppressor genes, we do not know whether the wild allele exerts a gain- or loss-of-function effect when compared to the S288C allele. We thus included both cases of increased or decreased suppressor gene copy number, as well as loss-of-function mutations. Indeed, we found that loss-of-function mutations in the essential query gene more frequently co-occurred with impactful mutations in their bypass suppressors than expected based on mutation frequency (**Fig. 4D**, **Data S8**). The occurrence of variants that can act as bypass suppressors may thus allow the accumulation of otherwise deleterious mutations in the bypassed essential genes.

### Mechanisms of bypass suppression by natural variants

We previously established a classification system to assign suppression interactions to distinct mechanistic categories on the basis of the functional relationship between the suppressor and query (suppressed) genes, such as annotation to the same biological process (Van Leeuwen et al. 2016). Using this classification system, we found that 22% of the bypass suppression interactions involving a natural variant could be explained by a functional relationship between the suppressor and the deleted essential gene (**Fig. 4E**). This fraction of functionally related gene pairs is significantly lower than that of essential gene bypass suppressors that were isolated through spontaneous mutation in the laboratory reference strain S288C (68%, *p* < 0.0005, Fisher’s exact test) (Van Leeuwen et al. 2020). The spontaneous suppressors isolated in the lab also frequently involved members of the same complex or pathway as the bypassed essential gene, but we did not find any such cases among the validated natural bypass suppressors (**Fig. 4E**, *p* < 0.05, Fisher’s exact test). Genes encoding members of the same complex or pathway as an essential gene are expected to be important for optimal cellular fitness themselves, making suppressor mutations that disrupt such genes less likely to occur in natural populations. Indeed, while 34% of the bypass suppressor mutations isolated in a laboratory setting occurred in an essential gene, only 9% of natural suppressors affected genes that are required for viability (**Fig. 4F**). Consistent with these observations, in seven out of ten cases where both natural variants and spontaneous laboratory mutations could bypass the same essential gene, the suppressor genes differed. Thus, natural bypass suppression variants tend to employ more indirect mechanisms than bypass suppressors isolated in a laboratory setting, likely because mutations affecting genes that are closely related to the bypassed gene are too deleterious to become fixed in natural populations.

### Loss of Msn5 suppresses *rad53*Δ lethality by restoring histone degradation

To gain further insight into the mechanisms underlying bypass suppression by natural variants, we investigated two examples in detail. The first example involves bypass of *RAD53*, which encodes a checkpoint kinase that in response to DNA damage or replication stress phosphorylates targets that induce a cell cycle delay, DNA repair, or replication fork stabilization (Allen et al. 1994, Weinert et al. 1994, Huang et al. 1998, Duncker et al. 2002, Lee et al. 2003). In our screen, *RAD53* was found to be essential in all tested genetic backgrounds except YJM975 (**Fig. 1D**). We identified a premature stop codon in *MSN5,* encoding a β-karyopherin that mediates the nuclear import and export of proteins and mature tRNAs (Alepuz et al. 1999, Yoshida and Blobel 2001, Murthi et al. 2010), as a potential suppressor variant (**Fig. 3A**, **4A**, **S4**). Indeed, deletion of *MSN5* rescued the lethality of *rad53*Δ in the S288C genetic background (**Fig. 5A**) and could also suppress the sensitivity to DNA damaging agents of *rad53-5004*, a partial loss-of-function allele that carries a premature stop-codon at residue S743 (**Fig. S7A**).

**Figure 5.**
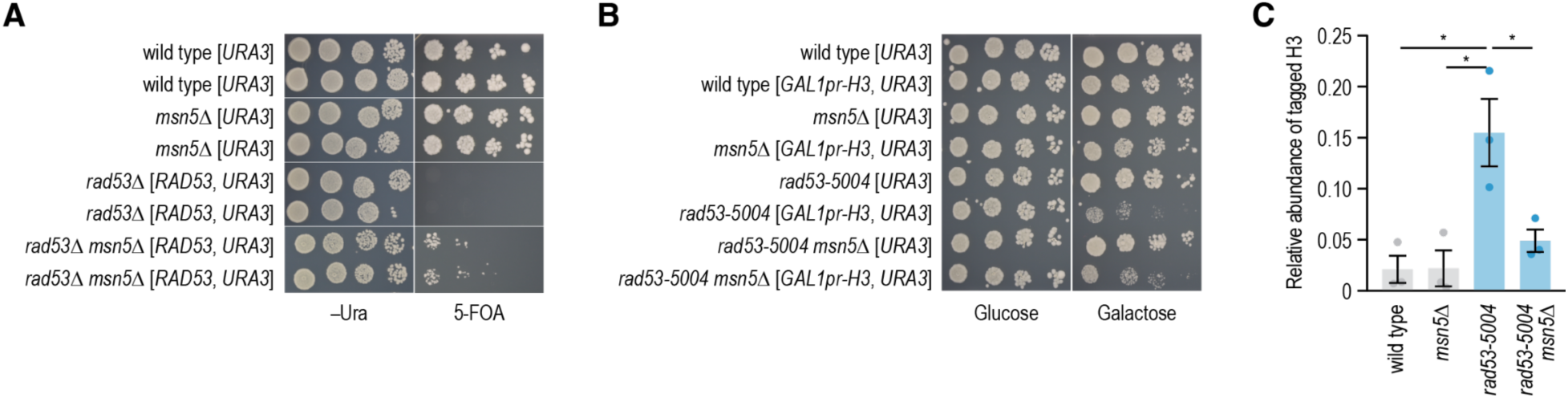
Loss-of-function mutations in *MSN5* suppress *rad53*Δ lethality by restoring histone degradation. (**A**) Loss of *MSN5* can suppress *rad53*Δ lethality. Cultures of the indicated strains were diluted to an optical density at 600 nm of 1, and a series of 10-fold dilutions was spotted on agar plates and incubated at 30°C for three days. (**B**) Deletion of *MSN5* reduces sensitivity of *rad53-5004* to histone overexpression. Spot dilutions as in (A) for wild-type, *msn5*Δ, *rad53-5004*, and *rad53-5004 msn5*Δ strains expressing a galactose inducible, His10-TEV-HA-tagged histone H3 gene (*GAL1pr-H3*) or the empty vector (Gunjan and Verreault 2003). (**C**) Histone degradation is restored in *rad53-5004 msn5*Δ double mutants. The strains described in (B) were cultured in media containing galactose to overexpress histone H3. Cells were then switched to media containing glucose and harvested after 80 min. The levels of tagged H3 were quantitated by Western blotting (**Fig. S7G**) and normalized to the total protein levels and to the amount of tagged H3 present at the time of glucose addition. Cells were blocked and kept in G1 using α-factor for the duration of the experiment. Error bars represent the standard error of the mean of three independent experiments. Statistical significance was determined using unpaired two-tailed t-tests, * *p* < 0.05.

S288C cells lacking *RAD53* attempt to replicate their DNA with suboptimal nucleotide levels, which together with an inappropriate entry into S phase leads to segregation of incompletely replicated chromosomes, causing lethality. Several *rad53*Δ bypass suppressors have been described in laboratory strains (Desany et al. 1998, Zhao et al. 1998, de Bruin et al. 2006, Garcia-Rodriguez et al. 2012, Manfrini et al. 2012). These suppressors either increase nucleotide pools or slow down the cell cycle, thereby allowing *rad53*Δ mutants to survive. However, *msn5*Δ neither delayed cell cycle progression nor affected nucleotide levels of a *rad53-5004* mutant (**Fig. S7B-D**). Furthermore, deletion of the G1 cyclin *CLN2*, which was previously shown to bypass the essential function of Rad53 by delaying entry into S-phase (Manfrini et al. 2012), further improved the fitness of a *rad53Δ msn5*Δ mutant both in the absence and presence of DNA damaging agent hydroxyurea (**Fig. S7E**). This suggests that suppression by *MSN5* occurs through a different mechanism than suppression by *CLN2*. Finally, cell cycle progression as well as the control of dNTP pools are regulated by both Rad53 and its upstream activator, the checkpoint kinase Mec1. However, loss of *MSN5* could suppress *rad53*Δ but not *mec1*Δ (**Fig. S7F**). Together, these results suggest that suppression by *MSN5* does not occur through increased nucleotide pools or decreased cell cycle progression.

Rad53, but not Mec1, is also required for the phosphorylation and subsequent degradation of excess histones that are not packaged into chromatin (Gunjan and Verreault 2003, Singh et al. 2009). In *rad53* mutants, excess histones bind non-specifically to the DNA, thereby adversely affecting DNA replication and repair. As a result, *rad53* mutants are highly sensitive to histone overexpression. To investigate whether *msn5*Δ could rescue this phenotype, we expressed histone H3 under control of a galactose-inducible promoter, leading to overexpression of histone H3 in media containing galactose (Gunjan and Verreault 2003). Lack of *MSN5* slightly but reproducibly rescued the growth defect of *rad53-5004* cells in the presence of galactose (**Fig. 5B**). Moreover, the defect in histone degradation of the *rad53-5004* mutant was suppressed in *rad53-5004 msn5*Δ double mutants (**Fig. 5C**, **S7G**). Loss-of-function mutations in *MSN5* may thus partially bypass *rad53*Δ by restoring histone degradation.

### Natural variants in *MKT1* bypass the need for mRNA decapping

In a second example, we investigated the bypass of *DCP1* and *DCP2* that were essential in S288C but not in most other genetic backgrounds (**Fig. 1D**). We further validated this finding by creating heterozygous deletion mutants of *DCP1* and *DCP2* in the genetic backgrounds S288C, IY_03-5-30-1-1-1_(1), and YPS163. Haploid progeny carrying the *dcp1*Δ or *dcp2*Δ alleles were indeed viable in the two natural isolates, but not in the reference strain S288C (**Fig. S8A**). Dcp1 and Dcp2 form the decapping complex that removes the 5′ 7-methylguanosine cap from mRNAs, thereby repressing translation of the mRNAs and targeting them for degradation by the exonuclease Xrn1 (Dunckley and Parker 1999). Xrn1 is not required for viability in S288C, suggesting that the essential function of Dcp1 and Dcp2 is to prevent over-translation of certain mRNAs through decapping, rather than enabling mRNA degradation.

The decapping complex is localized in processing bodies (P-bodies); granules consisting of mRNA and proteins, where translationally repressed mRNAs are either degraded or stored for later use (Wang et al. 2018). In unstressed S288C cells, P-bodies are small and few in number, but their formation gets quickly induced by osmotic stress (Shenton et al. 2006) (**Fig. 6A**). We found that compared to S288C, wild strains IY_03-5-30-1-1-1_(1) and YPS163 had low translation rates (**Fig. S8B**) and high levels of P-bodies (**Fig. 6B**) that did not further increase under stress conditions. Furthermore, Dcp1 and Dcp2 were required for P-body formation in S288C, but not in the two wild isolates (**Fig. 6A-B**).

**Figure 6.**
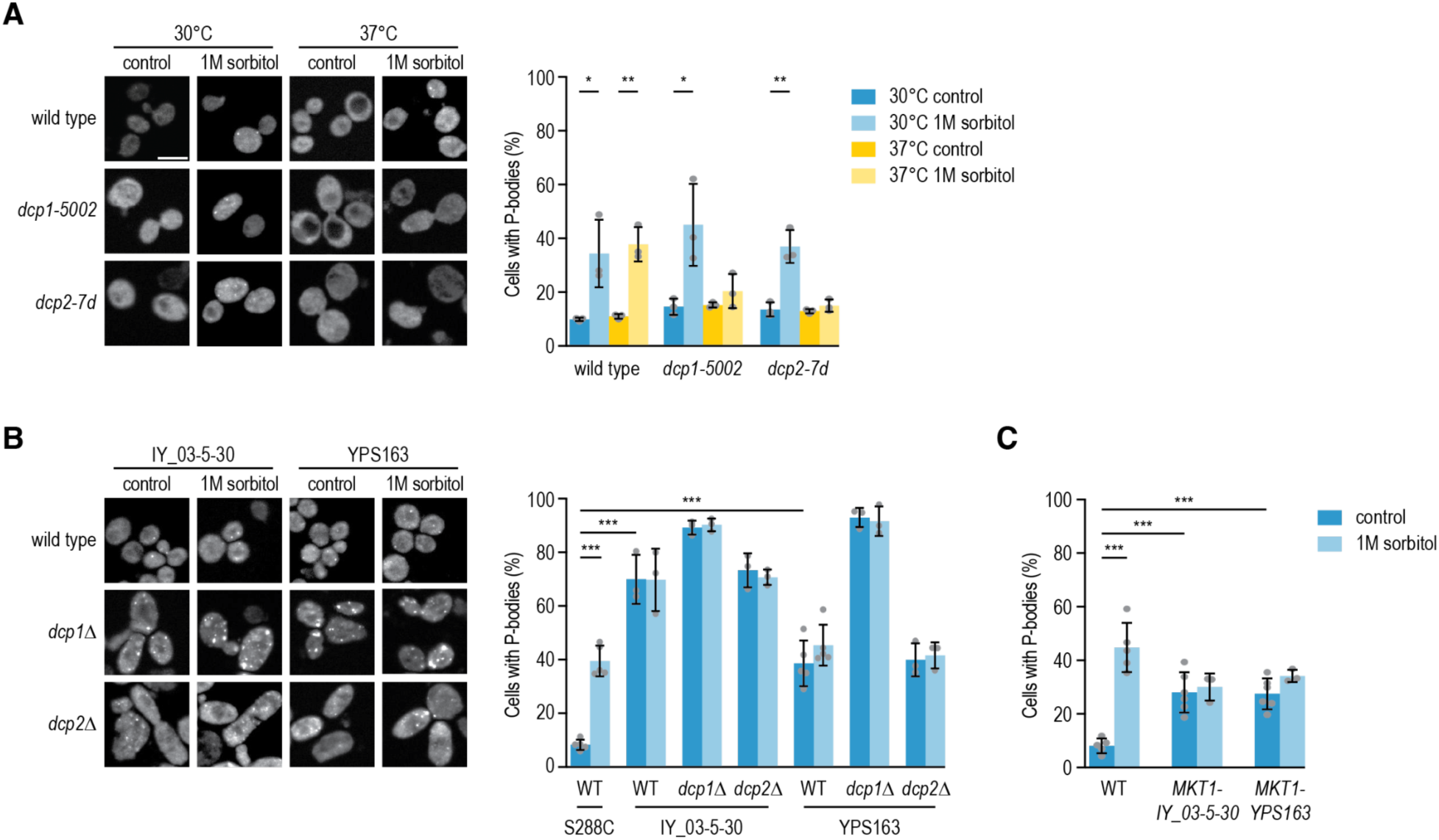
Natural variants in *MKT1* bypass the need for mRNA decapping. (**A**-**C**) P-bodies were visualized in the indicated strains using Dhh1-GFP. Shown are representative epifluorescent micrographs of exponentially growing cells, incubated in the absence or presence of 1 M sorbitol for 15 minutes. Measurements were performed in triplicate on at least 100 cells per replicate, and averages with standard deviations are shown. Strains in (A) and (C) have the S288C background. The alleles *dcp1-5002* and *dcp2-7d* are temperature sensitive mutants. Scale bar, 5 µm. Statistical significance was determined using unpaired two-tailed t-tests, * *p* < 0.05, ** *p* < 0.005, *** *p* < 0.0005. IY_03-5-30 = IY_03-5-30-1-1-1_(1).

We mapped the natural variants responsible for the bypass of the decapping complex in the wild strains to *MKT1* (**Fig. 3B**, **4A**, **S4**). Variants in *MKT1* are known to affect many phenotypes, and previous studies speculated that Mkt1’s pleiotropic effect could result from it altering expression of many genes, possibly through regulating mRNA storage in P-bodies (Lee et al. 2009, Lewis et al. 2014). Indeed, when we introduced the wild alleles of *MKT1* into the S288C background, we observed high levels of P-bodies under standard growth conditions, consistent with the phenotype observed in the natural isolates (**Fig. 6B-C**). Together, these data suggest that absence of mRNA decapping can be compensated for by increased mRNA storage in P-bodies in an Mkt1-dependent manner, thereby maintaining a balanced pool of mRNAs that are available for translation.

## DISCUSSION

We systematically tested the essentiality of 763 budding yeast genes across 18 genetically diverse backgrounds. By measuring the consequence of deleting each gene individually, including an average of 24 replicates (six biological and four technical replicates) per gene, and validating all cases of context-dependent essentiality in a secondary assay, we obtained high experimental accuracy that allowed us to confidently classify genes as either essential or nonessential in a genetic background. We discovered 39 genes that were essential in S288C but not in one or more of the natural isolates (**Fig. 1D**), mapped the genomic loci responsible for this difference for 37 cases (**Fig. 3B**), and identified 24 of the underlying suppressor genes (**Fig. 4A**).

### Prevalence of bypass suppression

Around 5% of the tested genes showed context-dependent gene essentiality in our screen. Similarly, previous studies reported that 1-4% of genes classified as essential in the laboratory strain S288C showed differential essentiality across natural budding yeast isolates (Dowell et al. 2010, Chen et al. 2022). By contrast, over three times as many essential genes (∼17%) can be bypassed by spontaneous mutations during experimental evolution in a laboratory (Van Leeuwen et al. 2020, Couce et al. 2024). Suppressor variants are often deleterious when they occur in the absence of the suppressed query mutation (Ünlü et al. 2023) and the spontaneous variants isolated in a laboratory may thus not be present in wild genomes, resulting in fewer essential genes being bypassed by natural variants than in laboratory evolution assays. Nonetheless, bypass suppression by natural variants was relatively common and multiple instances were found in every tested wild strain.

Because of the stringent threshold we applied for calling a gene nonessential in our secondary validation assay (**Fig. 1E-F**, **Data S3**), we may have missed cases of context-dependent essentiality that required the presence of four or more modifier loci. However, although our threshold would have allowed us to identify cases that required simultaneous mutation of two or three suppressor alleles at different loci, we were not able to validate any examples (**Fig. S5B**), and all 24 validated natural bypass suppressors acted in isolation. Moreover, our screen identified 80% of the genes that were previously shown to be essential in S288C but nonessential in Σ1278b using a different method (Dowell et al. 2010), confirming that our assay detected most cases of context-dependent gene essentiality.

### Complexity of bypass suppression

Although multiple studies have previously concluded that context-dependent gene essentiality often has complex underpinnings in yeast (Dowell et al. 2010, Hou et al. 2019), we find that differences in gene essentiality are often driven by a single modifier. While these findings appear contradictory, they are not necessarily in conflict with each other. We often detected multiple loci with selection for wild sequence in a cross (**Fig. 3**), suggesting complexity of the phenotype, but when validating the suppressor genes, we found that one locus is often sufficient for the change in gene essentiality (**Fig. S4**). This genetic simplicity is consistent with other studies describing a single gene or genomic locus driving a phenotypic trait (Johnston et al. 2011, Barson et al. 2015, Jones et al. 2018, Thompson et al. 2020, Parts et al. 2021). The additional loci that are not required for the change in essentiality may contribute to the overall fitness of the strain (**Fig. S5A**). Thus, the viability component of the growth phenotype is often genetically “simple”, but the fitness component can be complex. We suspect that, compared to the many ways in which the fitness of a strain can be improved (Parts et al. 2021), there are relatively few ways to bypass the requirement for an essential gene by standing variation, and thus few variants are expected to contribute to this phenotype within a single genetic background.

### Mechanisms of bypass suppression

The natural bypass suppressors tended to be different from other types of suppressors that had been described previously. In cases where both natural (this study) and laboratory (Van Leeuwen et al. 2020) bypass suppressors had been identified for the same essential gene, the suppressor genes were different in 70% of the cases. In general, natural suppressors more indirectly bypassed the need for an essential gene compared to those isolated in a laboratory (**Fig. 4E**), for example by altering the pool of mRNA available for translation. Furthermore, natural suppressors tended to be more diverse than laboratory ones, and context-dependent essential genes that were nonessential in multiple wild strains were often bypassed by different suppressor genes in each background (**Fig. 3B**). These results are consistent with and extend the previous finding that yeast gene essentiality switched multiple times during evolution (Chen et al. 2022).

We previously identified natural variants that could suppress the fitness defect at high temperature of essential gene temperature sensitive alleles (Parts et al. 2021). These suppressors were dependent on the presence of the temperature sensitive allele and thus did not necessarily reveal differences in gene essentiality between wild isolates. Comparison of essential gene deletion and temperature sensitive mutant suppressors showed little overlap in the suppressible genes (**Fig. 4C**). This is in agreement with our previous finding that suppressors of TS alleles frequently affect stability or expression of the TS mutant by perturbing mRNA or protein turnover (Van Leeuwen et al. 2016), as such suppressors are not expected to rescue deletion alleles. Nonetheless, variants in *MKT1* could suppress the phenotype of many diverse mutant alleles, including both TS and deletion mutants (**Fig. 3B**, **4A**) (Johnson et al. 2019, Parts et al. 2021, Hale et al. 2024). Here, we showed that wild alleles of *MKT1* increase P-body formation (**Fig. 6C**), thereby likely affecting the expression of a large part of the genome, and explaining its pleiotropic effects on many phenotypes and mutants (Fay 2013, Albert et al. 2014, Lewis et al. 2014, Parts et al. 2014, Albert et al. 2018).

Around 88% of natural bypass suppression interactions (21 out of 24) revealed new connections between genes. Among the newly identified bypass suppressors was *MSN5*, which could partially bypass *rad53*Δ by restoring histone degradation (**Fig. 5**). The exact role of β-karyopherin Msn5 in the degradation of excess histones remains unclear. Msn5 is involved in the nuclear import and export of proteins and mature tRNAs (Alepuz et al. 1999, Yoshida and Blobel 2001, Murthi et al. 2010). Histones are not known to be transported by Msn5; the karyopherins Kap95, Kap114, Kap123 and Pse1 (Kap121) are thought to be the main importers of core histones into the nucleus (Mosammaparast et al. 2001, Mosammaparast et al. 2002). Possibly, the levels of these importers are changed in an *msn5*Δ mutant, thereby affecting histone import. Alternatively, Msn5 may play a role in the transport of linker histone H1 or of the histone chaperone Nap1. Further research will be needed to distinguish between these possibilities.

### Future perspectives

Overall, our results provide insight into the prevalence, complexity, and mechanisms of bypass suppression by standing variation. We find that bypass suppression is relatively common: we uncovered on average nine examples of bypass suppression per natural yeast strain. The mapped suppression interactions were also relevant for natural budding yeast populations, in which naturally occurring loss-of-function mutations in the essential query gene tended to be accompanied by impactful mutations in the corresponding bypass suppressor gene (**Fig. 4D**) (Pons and Van Leeuwen 2023). Around 25% of essential genes in any given human cell line can be classified as context-specific essential (Hart et al. 2015), suggesting that bypass suppression may also be relatively common in the genomes of more complex cells. A thorough understanding of how modifier variants influence relatively simple phenotypes, such as gene dispensability, will be crucial for the interpretation of more complex cases and will refine our view of variable penetrance across phenotypes, species, and contexts, including human disease.

## METHODS

### Yeast strains and growth conditions

Yeast strains were grown using standard rich (YPD) or minimal (SD) media. To identify genetic context-dependent essential genes (see next section), we used strains from the TS-allele-on-plasmid collection (*MATα xxx*Δ*::KlLeu2_natR(Cterm) his3*Δ*1 leu2*Δ*0 ura3*Δ*0* [*xxx-ts_natR(Nterm)*, *AgSTE3pr-hphR*, *URA3*] and *MAT**a** xxx*Δ*::natR_kanR(Cterm) his3*Δ*1 leu2*Δ*0 ura3*Δ*0* [*xxx-ts_kanR(Nterm)*, *AgMFA2pr-hphR*, *URA3*]) (Van Leeuwen et al. 2020). For the allele swaps (see section “suppressor candidate validation”), we used strains from the BY4741 and BY4742 deletion mutant collections (*MAT**a** xxx*Δ*::kanMX4 his3*Δ*1 leu2*Δ*0 ura3*Δ*0 met15*Δ*0* and *MATα xxx*Δ*::kanMX4 his3*Δ*1 leu2*Δ*0 ura3*Δ*0 lys2*Δ*0*; Euroscarf). All other yeast strains used in this study are listed in **Data S9**.

### Saccharomyces cerevisiae phylogeny

We selected 127 *Saccharomyces cerevisiae* isolates representative of the main lineages from the 1,011 (Peter et al. 2018) and the Chinese (Duan et al. 2018) *Saccharomyces cerevisiae* collections (**Data S10**). Reads were mapped against the *Saccharomyces cerevisiae* reference genome (S288C, version R64.3.1) using BWA-MEM and indexed with BWA v0.7.17 (Li and Durbin 2009). The resulting SAM file was filtered for PCR and optical duplicate reads as well as unmapped reads using SAMtools v1.19 *view* (options: -bS -F 1036) (Li et al. 2009) and converted into the binary alignment map (BAM) format. The BAM file was sorted and indexed with SAMtools *sort* and SAMtools *index*. We extracted whole-genome genotypes with SAMtools *mpileup* (options: -u -min-MQ5 --output-tags AD,ADF,ADR,DP,SP --skip-indels --redo-BAQ) and BCFtools v1.19 *call* (options: -c -Oz) and generated a gVCF for each isolate. Then, we filtered the gVCF files using VCFtools v0.1.17 (Danecek et al. 2011), applying criteria for depth (--min-DP 10) and call quality (--min-Q 20). Only biallelic positions were retained (option: --max-alleles 2 and --min-alleles 2), while indels and mtDNA were removed (--remove-indels; custom command: *grep -v* chrMT). The filtered gVCF files were zipped with bgzip v1.16 and merged into a single gVCF file with BCFtools *merge*. We extracted the SNPs with PLINK v1.90b7 (Purcell et al. 2007) (--snps-only ‘just-acgt’) and filtered out those with missing call rates exceeding 0 (--geno 0), not allowing for positions with missing genotype information.

For the phylogeny construction (**Fig. 1A**, **S2C**), the derived VCF file was converted into a PHYLIP file, a file that stores a multiple sequence alignment of DNA sequences obtained concatenating the bases at the SNP positions for each isolate (vcf2phylip.py v.2.8) (Ortiz 2019). The phylogenetic tree was generated with IQ-TREE v1.6.12) which automatically selected the best-fit model TVMe+R2 (options: -bb 1000 -alrt 1000 -bnni -nt AUTO; for reproducibility seed: 235075) (Nguyen et al. 2015, Kalyaanamoorthy et al. 2017, Hoang et al. 2018). The .*treefile* in newick format was depicted in an unrooted topology and further edited with the Interactive Tree Of Life v5 (Letunic and Bork 2021).

### Identifying genetic context-dependent essential genes

To identify genes that were essential in S288C but nonessential in other genetic contexts, the TS-allele-on-plasmid collection was used to cross essential gene deletion alleles into wild yeast isolates by SGA analysis (**Fig. 1B**). The TS-allele-on-plasmid collection consists of haploid strains, each deleted for an essential gene in their genome but viable because of the presence of a TS allele of the same essential gene on a counter-selectable plasmid. Because this collection contains TS alleles of the essential genes, all experiments with strains carrying the plasmid were performed at 26°C to ensure proper functioning of the encoded essential protein.

#### Making the TS-allele-on-plasmid collection SGA compatible

To be able to introduce the essential gene deletion alleles into the wild strains by SGA, the SGA markers (*can1*Δ*::STE2pr-SpHIS5* and *lyp1*Δ) that are used to select haploid cells had to be introduced into the TS-allele-on-plasmid collection. The TS-allele-on-plasmid collection contains strains with two different genotypes: *MATα xxx*Δ*::KlLeu2_natR(Cterm) his3*Δ*1 leu2*Δ*0 ura3*Δ*0* [*xxx-ts_natR(Nterm)*, *AgSTE3pr-hphR*, *URA3*] and *MAT**a** xxx*Δ*::natR_kanR(Cterm) his3*Δ*1 leu2*Δ*0 ura3*Δ*0* [*xxx-ts_kanR(Nterm)*, *AgMFA2pr-hphR*, *URA3*]. SGA markers were introduced into the TS-allele-on-plasmid strains using SGA analysis (Baryshnikova et al. 2010). In brief, the *MATα* TS-allele-on-plasmid strains were crossed to LY00044 (*MAT**a** ho*Δ*::natMX4 can1*Δ*::STE2pr-SpHIS5 lyp1*Δ*0 his3*Δ*1 leu2*Δ*0 ura3*Δ*0 met15*Δ*0*; **Data S9**). In a series of subsequent pinning steps, diploid cells were selected and sporulated, haploid cells were selected, and finally SD(MSG) –Arg/Lys/Ura/Leu +CAN/LYP/HYG/NAT was used to isolate *MATα* strains carrying the essential gene deletion allele, the plasmid carrying a TS allele of the essential gene, and the SGA markers. The final genotype of the isolated strains was MAT*α xxx*Δ*::KlLEU2_natR(Cterm) can1*Δ*::STE2pr-SpHIS5 lyp1*Δ*0 ho*Δ*::natMX4 his3*Δ*1 leu2*Δ*0 ura3*Δ*0* [*xxx-ts_natR(Nterm)*, *AgSTE3pr-hphR*, *URA3*].

Similarly, to introduce the SGA markers into the *MAT**a*** TS-allele-on-plasmid strains, they were crossed to LY00087 (*MATα can1*Δ*::SpHIS5 lyp1*Δ*::STE3pr-LEU2 his3*Δ*1 leu2*Δ*0 ura3*Δ*0 met15*Δ*0*; **Data S9**). In a series of subsequent pinning steps, diploid cells were selected and sporulated, haploid cells were selected, and finally SD(MSG) –Arg/Lys/Ura/His/Leu +CAN/LYP/NAT was used to isolate *MATα* strains carrying the essential gene deletion allele, the plasmid carrying a TS allele of the essential gene, and the SGA markers. The final genotype of the isolated strains was *MATα xxx*Δ*::natR_kanR(Cterm) can1*Δ*::SpHIS5 lyp1*Δ*::STE3pr-LEU2 his3*Δ*1 leu2*Δ*0 ura3*Δ*0* [*xxx-ts_kanR(Nterm)*, *AgMFA2pr-hphR*, *URA3*].

#### Making the wild yeast isolates SGA compatible

We previously constructed 11 wild isolates that were compatible with SGA analysis by deleting auxotrophic marker genes from the genome (*MAT**a** ho*Δ*::hphMX6 ura3*Δ*::kanMX4 his3*Δ*1 leu2*Δ*0*; **Data S9**) (Parts et al. 2021). To increase the genetic diversity in our strain collection, we constructed another eight SGA-compatible wild strains (**Fig. 1A**). Previously, Fournier *et al*. deleted *HO* in a set of 55 wild yeast strains and isolated haploid spores (*MAT**a** ho*Δ*::kanMX*) (Fournier et al. 2019). We selected a subset of eight strains that we further deleted for *URA3*, *HIS3*, and *LEU2*. First, guide RNA (gRNA) sequences targeting *URA3*, *HIS3*, or *LEU2* were cloned into the pML104-HygMX4 (for *URA3*) or pML104 (for *HIS3* and *LEU2*) plasmid vectors, which carry Cas9 and either *hphR* or *URA3* (**Data S9**) (Laughery et al. 2015). Second, we amplified the *ura3*Δ*0*, *his3*Δ*1*, and *leu2*Δ*0* deletion cassettes from BY4741 (*MAT**a** his3*Δ*1 leu2*Δ*0 ura3*Δ*0 met15*Δ*0*; **Data S9**) by PCR, thereby including ∼400 bp upstream and downstream of the open reading frame, and co-transformed the PCR fragment and the corresponding gRNA plasmid into the wild strains. The gRNAs will cut the auxotrophic marker gene and the homology of the promoter and terminator sequences of the PCR product to the genomic sequences flanking the double-stranded DNA breaks will promote repair via homologous recombination and integration of the PCR product into the genome. Transformants were initially isolated on media selecting for the gRNA plasmid and then propagated on YPD. Within three days of growth on YPD, the vast majority of yeast strains had lost the gRNA plasmids and had properly replaced the auxotrophic marker gene with the (partial) deletion allele, which we confirmed by testing for growth on the appropriate selective media, PCR, Sanger sequencing, and whole genome sequencing. During sequencing, WI00236 was found to contain a transposon insertion in *URA3*, rather than the intended deletion allele. As *URA3* was still inactivated, we continued to use this strain for further experiments. The final genotype of the strains was *MAT**a** ho*Δ*::kanMX his3*Δ*1 leu2*Δ*0 ura3::Ty* for WI00236 and *MAT**a** ho*Δ*::kanMX his3*Δ*1 leu2*Δ*0 ura3*Δ*0* for all other wild isolates (**Data S9**).

#### SGA analysis and scoring

Next, the SGA-compatible TS-allele-on-plasmid collection and wild strains were crossed using SGA analysis (**Fig. 1B**). SGA analysis was performed as described previously (Baryshnikova et al. 2010), with the exception that 5% mannose was added to the YPD plates used in the first steps of SGA to facilitate pinning of the wild isolates. In brief, the 18 SGA-compatible *kanMX*-marked wild strains (**Data S9**; *MAT**a** ho*Δ*::hphMX6 ura3*Δ*::kanMX4 his3*Δ*1 leu2*Δ*0* or *MAT**a** ho*Δ*::kanMX his3*Δ*1 leu2*Δ*0 ura3*Δ*0*), and an S288C negative control strain (LY00045, *MAT**a** ura3*Δ*::kanMX4 his3*Δ*1 leu2*Δ*0 met15*Δ*0* or DMA809, *MAT**a** ho*Δ*::kanMX4 his3*Δ*1 leu2*Δ*0 ura3*Δ*0 met15*Δ*0*; **Data S9**) were crossed to the TS-allele-on-plasmid collection carrying the SGA markers. Each cross was performed in four technical replicates. In a series of subsequent pinning steps, diploid cells were selected and sporulated, and colonies consisting of pools of around 60,000 haploid segregant progeny (Parts et al. 2014) carrying both the essential gene deletion allele and the plasmid carrying a TS allele of the essential gene were isolated. Finally, 5-fluoroorotic acid (5-FOA) was used to remove the plasmid, resulting in loss of the essential gene. Plates were imaged and colony size was measured as pixel area (**Data S2**) (Wagih and Parts 2014). The screen was repeated for all TS-allele-on-plasmid strains that appeared to grow in the presence of 5-FOA in at least one of the wild strain crosses. UWOPS87-2421 (LY00011) carried an aneuploidy of chromosome IV in the initial screen, but not in the validation screen, and thus only colony size values from the validation screen were used for genes located on chromosome IV for this strain.

To correct for differences in fitness between the natural isolates, raw colony size measurements were normalized using the median colony size of wild strain progeny before 5-FOA selection (**Data S2**). Spontaneous mutations in *URA3* can sometimes occur in one of the technical replicates of a cross and cause resistance to 5-FOA. To remove such cases from consideration, we used Grubbs’ test to identify and filter outlier colonies. Finally, we compared the fitness of the haploid progeny in the presence of 5-FOA between the S288C control cross and the wild strain crosses, and performed a one-sided Welch’s t-test to determine statistical significance of observed differences. Mutants that had a mean normalized fitness > 0.25 in the presence of 5-FOA in the S288C control cross were considered to be alive and excluded. Genes were considered to be nonessential in progeny of a wild strain cross if more than 50% of the strains deleted for the gene had a mean normalized fitness that was at least 0.25 higher in the wild strain progeny than in the corresponding S288C control progeny, and the associated *p* value was < 0.02. Genes located on the chromosome VIII-XVI translocation or on the chromosome XIV duplication (**Fig. 1C**, **S1A-B**) were not further considered. In total, 124 genes that were essential in S288C appeared to be nonessential in at least one of the wild strain crosses (**Data S2**).

#### Validation by random sporulation analysis

To validate context-dependent essentiality, SGA-compatible TS-allele-on-plasmid strains of all 124 identified context-dependent essential gene candidates were crossed to the relevant SGA-compatible wild isolates and sporulated. As a control, we crossed each essential gene mutant to the reference strain S288C (LY00045; **Data S9**) as well. Sporulated cells were plated in parallel onto two agar plates that selected for haploid spores that carried the essential gene deletion allele: one plate with media that also selected for the plasmid carrying the TS allele (SD(MSG) – Arg/Lys/His/Ura/Leu +CAN/LYP/G418 or SD(MSG) –Arg/Lys/Ura/Leu +CAN/LYP/G418/NAT) and the second plate with media that selected against the plasmid (SD(MSG) –Arg/Lys/His/Leu +CAN/LYP/G418/5-FOA or SD(MSG) –Arg/Lys/Leu +CAN/LYP/G418/NAT/5-FOA). After four days at 30°C or five days at 26°C, plates were imaged (**Fig. 1E-F**), and colony number was determined using Cell Profiler (Carpenter et al. 2006). A gene was considered to be nonessential in a genetic background when at least 10% of the colonies that grew on –Ura media could also grow on media with 5-FOA (**Fig. 1D**, **Data S3**, **S4**). For two genes, *TFG2* and *NUP159*, the experiment failed for technical reasons and these genes were not further considered.

### Comparison to other datasets

For comparison of context-dependent essential genes identified here to similar gene sets identified previously (**Fig. 2B**, **S2A**), we used the following gene sets: “Polymorphic genes” (Chen et al. 2022), “Background healthy genes” (Caudal et al. 2022), “Strain-specific genes” (Wang et al. 2022), and “Dispensable essential genes” (Van Leeuwen et al. 2020). For each study, the fold enrichment was calculated as the fraction of context-dependent essential genes identified by us that also showed context-dependency in the other study, divided by the fraction of genes that were always essential in our screen that did show context-dependency in the other study. Only genes that were included in both our screen and in the study of interest were considered. Statistical significance of the overlap was assessed by Fisher’s exact tests.

To investigate whether genes were mutated in other natural yeast strains, we used data from the 1,011 genomes project (Peter et al. 2018). For the analyses in **Fig. 2A** and **S6B**, we only considered homozygous loss-of-function mutations in genes. For the analysis in **Fig. 4D**, we considered either homozygous loss-of-function mutations or absence for the context-dependent essential gene and absence, copy number increase, or homozygous loss-of-function mutations for the suppressor gene. In all cases, we used mutation annotations from the 1,011 genomes project (http://1002genomes.u-strasbg.fr/files). Strains that were used in our screen were excluded from the analysis.

For the comparison of context-dependent essential genes to genes that have a temperature sensitive (TS) allele that can be suppressed by a natural variant (**Fig. 4C**), we used suppression values from (Parts et al. 2021). These suppression values were calculated as the difference in fitness between haploid segregants carrying a TS allele that were derived from an isogenic S288C cross or from a cross between S288C and a wild strain. TS alleles that were not temperature sensitive in the S288C cross and values that were affected by a translocation or an aneuploidy were removed from consideration. TS mutants that had a suppression value > 0.75 in a given background were considered suppressed. If a gene was represented by multiple TS alleles, we considered a gene suppressed in a background as soon as one of the alleles was suppressed.

### Properties of dispensable essential genes

For the analysis of enrichment of context-dependent essential genes for other gene- or protein-level properties (**Fig. S2B**), we compared feature values of context-dependent essential genes to those of genes that were essential in all tested yeast strains. We considered gene copy number (i.e., number of paralogs) (Koch et al. 2012), gene length, transcript count (i.e., expression level) (Lipson et al. 2009), expression variation under different environmental conditions (Gasch et al. 2000), coexpression degree calculated as the number of genes with similar expression profiles (i.e., MEFIT scores > 1) (Huttenhower et al. 2006), multifunctionality calculated as the number of GO SLIM biological process annotations (Ashburner et al. 2000), number of species in which the gene is conserved (Koch et al. 2012), cocomplex degree (the number of proteins that share a complex with a protein of interest), and whether the gene encoded a membrane-associated protein (Babu et al. 2012) or a member of a protein complex (Meldal et al. 2021). For members of protein complexes, we counted the number of distinct complexes in which they were found and calculated their cocomplex degree as the number of protein partners present in those complexes.

### Classifying essential genes as either context-dependent or core essential

To define a core set of essential genes (**Data S5**), we used five sources of context-dependent essential genes. First, we used Dataset EV13 from (Van Leeuwen et al. 2020), that classified essential genes as either “core” or “dispensable” essential, based on whether spontaneous bypass suppressors could be identified that bypassed the requirement for the essential gene in reference strain S288C. We considered all “dispensable” essential genes as context-dependent essential. Dataset EV13 also contained a "Not further classified" category for genes that were essential in S288C and nonessential in another genetic background. For the current study, we considered those as context-dependent essential. Second, we used Dataset S7 from (Chen et al. 2022), that listed genes “exhibiting essentiality polymorphism” based on the effect of gene inactivation on viability in 15 yeast isolates. We classified these genes as context-dependent essential. Third, we used data from (Dowell et al. 2010), who tested the viability of gene deletion mutants in Σ1278b. We considered genes that were essential for viability in S288C but not in Σ1278b to be context-dependent essential. Fourth, we used the phenotype of genes in our screen (**Data S4**). Finally, we used mutation annotations from the 1,011 genomes project (http://1002genomes.u-strasbg.fr/files) to identify genes that carried homozygous loss-of-function mutations in one or more yeast isolates. These genes were considered “potential context-dependent essential”, as it is possible that loss-of-function mutations that occur relatively late in the coding sequence produce a truncated but functional protein.

For the final classification, we considered a gene to be core essential if it was identified as essential in all four experimental datasets and did not carry homozygous loss-of-function mutations in any of the 1,011 yeast isolates. A gene was considered to be context-dependent essential if it was identified as such in at least one of the four experimental datasets. The remaining genes, that were not identified as context-dependent essential in one of the experimental studies but did carry homozygous loss-of-function mutations in at least one wild yeast strain, were not further classified.

### Sequencing, read mapping, and SNP calling

All wild strains were sequenced on the BGI DNBseq platform using paired-end 100-bp reads, with an average read depth of ∼90x. Reads were aligned to the S288C reference genome version R64.2.1 using BWA v0.7.17 (Li and Durbin 2009). Pileups were processed and variants were called using SAMtools/BCFtools v1.11 (Li et al. 2009). Variants that had a Phred quality score below 20 in all sequenced strains were removed from consideration. The consequence of detected variants was determined using Ensembl’s VEP (McLaren et al. 2016). All whole-genome sequencing data are publicly available at NCBI’s Sequence Read Archive (http://www.ncbi.nlm.nih.gov/sra), under accession number PRJNA1131954. Variants are listed in **Data S1**.

### QTL analysis

We performed bulk segregant analysis on 44 crosses that showed context-dependent gene essentiality. We collected ∼1,000 haploid progeny colonies for each cross, either in the presence or absence of the plasmid carrying the essential gene, using the random sporulation assay described above. Colonies were scraped from the agar plates, and genomic DNA was isolated from the pools using the Qiagen DNeasy Blood & Tissue kit. Samples were sequenced and SNP calling was performed as outlined in the previous section, with the exception that variants were called using the BCFtools option --ploidy 2 (even though the sequenced samples were haploid) to detect variants at a lenient cut-off. DP4 values in the resulting VCF file were used to calculate reference allele frequencies, thereby excluding variants that were not found in one of the parental strains. Resulting raw variant frequencies are listed in **Data S6**.

Qualimap v2.3 (Okonechnikov et al. 2016) was used to detect aneuploidies based on variation in sequencing read depth across chromosomes (**Data S11**). Aneuploidies were detected in five crosses: TSP389 x WI00237 (chrI), TSP2180 x WI00237 (chrIX), TSP2413 x WI00231 (chrI), TSP2425 x WI00235 (chrXI), and TSP3539 x WI00236 (chrI). In all cases, an extra copy of the S288C chromosome was present in both the control (with plasmid carrying the essential gene) and the 5-FOA (without plasmid) sample, suggesting that the aneuploidy was present in the S288C mutant strain that was used in the cross. None of the aneuploidies significantly changed in abundance between the 5-FOA and control samples, suggesting that they did not influence survival in the absence of the essential gene.

To detect QTLs (**Fig. 3**, **S3**), raw allele frequencies for each site in each cross were averaged in a 30kb window centered on the site. Outlier sites that had raw allele frequencies deviating by at least 0.25 from the window average, likely reflecting mapping issues, were filtered out, and the averaging performed again. Sites for which less than 15 SNPs were present in the 30kb window were filtered out. QTLs were called as regions of at least 0.2 reference allele frequency change between the control (with plasmid expressing the essential gene) and 5-FOA (without plasmid) conditions, with QTL location assigned to the site of the largest change in the region. QTLs mapping to selection markers used in the cross, to chromosome III or the mitochondrial DNA, or within 20kb of the subtelomere, were filtered out. A null distribution of expected number of QTLs between crosses was constructed from contrasting data from all possible pairs of control crosses of the same wild strain, using the same QTL calling approach. Detected QTLs are listed in **Data S7**.

### Gene knockout

To test whether knockout of an essential gene was lethal in a particular strain background (**Fig. S3B**), we used a previously described CRISPR-Cas9 based method (Laughery et al. 2015). In brief, we cloned gRNAs targeting the essential genes into a vector carrying Cas9 (pML107; **Data S9**). The designed gRNAs targeted regions in the genome that were identical in sequence in the wild and reference genomes. Strains were then transformed with the CRISPR plasmid as well as an 80 base pair double-stranded DNA repair template that introduced a single base pair deletion as well as two substitutions within the PAM sequence in the targeted gene.

Transformants were selected on SD –Leu media that selects for the plasmid. Due to the high efficiency of CRISPR-Cas9 mediated genome editing, no viable colonies are expected to be obtained when the targeted gene is essential for viability in a given background. To control for variation in transformation and genome editing efficiency between strains, each strain was also transformed with similar reagents targeting the nonessential gene *YFR054C*, and the number of viable colonies obtained for the essential gene knockout was calculated as a percentage of those obtained for *YFR054C* knockout. NCYC110 was excluded from this analysis, because of a low transformation and/or genome editing efficiency compared to the other strains.

### Suppressor candidate validation

To validate bypass suppressor candidates (**Fig. S4**, **S5**), we introduced potential suppressor alleles into the reference genetic background. We amplified the genes including ∼400-1,000 bp upstream of the start codon and ∼200-400 bp downstream of the stop codon from the various wild strains by PCR. The PCR product and a plasmid carrying Cas9 and a gRNA targeting kanR (pML104-kanR1136 and/or pML107-kanR468; **Data S9**) were co-transformed into a haploid strain carrying a deletion allele of the suppressor gene marked with kanMX4. The gRNA will cut the kanMX4 cassette and the homology of the promoter and terminator sequences of the PCR product to the genomic sequences flanking the double-stranded DNA breaks will promote repair via homologous recombination and integration of the wild allele into the genome. For essential genes, we used a similar strategy using strains from the TS-allele-on-plasmid collection (see “Yeast strains and growth conditions”) and the plasmids pML104-natR412 and/or pML107-natR854 (**Data S9**) that carry gRNAs that target natR. Transformants were initially selected on media selecting for the gRNA plasmid and then propagated on YPD. Within three days of growth on YPD, the vast majority of yeast strains had lost the gRNA plasmids and properly replaced the suppressor candidate deletion allele with the wild allele, which we confirmed by PCR and Sanger sequencing. For essential genes, we streaked the allele-swapped strains on SDall +5-FOA to remove the plasmid carrying the essential gene. For suppression of *srp68*Δ and *srp72*Δ by the Σ1278b allele of *UBR2*, we used an *ubr2*Δ mutant as the Σ1278b allele carries a premature stop codon and is thus thought to be a loss-of-function allele. For *rad53*Δ, the YJM975 and S288C alleles of *MSN5* were cloned into pRS313 and expressed over an *msn5*Δ allele (**Data S9**). Finally, for *ino80*Δ *MKT1*, *pfy1*Δ *IRA1*, *srp68*Δ *GET3*, and *srp72*Δ *GET3*, the alleles that were used in the allele swaps did not come from the wild strain than that was used in the QTL analysis, but from a different strain that carried similar nonsynonymous variants in the swapped gene (**Data S9**).

Next, we crossed the strains carrying a wild allele of the suppressor candidate to the corresponding TS-allele-on-plasmid mutant and sporulated the resulting diploids. We isolated haploid progeny carrying the TS-allele plasmid, the essential gene deletion allele, and the suppressor allele from the wild background. Growth of the isolated strains was tested in the presence of 5-FOA to determine whether the strains could survive without the essential gene.

### Histone H3 assays

To determine the effect of deletion of *MSN5* on the histone degradation defect of *rad53* mutants, we transformed the single and double mutant strains, as well as wild-type cells, with a plasmid expressing a galactose inducible, His10-TEV-HA-tagged histone H3 gene or empty vector (Gunjan and Verreault 2003) (**Data S9**). Because *rad53*Δ is lethal, the deletion allele was complemented with a *HIS3*-plasmid carrying the partial loss-of-function allele *rad53-5004* (**Data S9**). Transformants were grown overnight in synthetic media lacking histidine and uracil to select for both plasmids and supplemented with 2% raffinose. For spot dilution assays (**Fig. 5B**), cells were diluted to an optical density at 600 nm of 1, and a series of 10-fold dilutions was spotted on agar plates containing synthetic media lacking histidine and uracil supplemented with 2% glucose or 2% galactose and incubated at 30°C for three days. For Western Blotting (**Fig. 5C**, **S7G**), cells were diluted to an optical density at 600 nm of 0.5 in synthetic media lacking histidine and uracil for 2 hours and synchronized in G1 with α-factor for 1.5 hours. Histone H3 overexpression was induced by addition of 2% galactose while cells were maintained in G1 with α-factor. After 1.5 hours, cells were washed and resuspended in synthetic media lacking histidine and uracil supplemented with 2% glucose and α-factor. Cells were harvested before galactose addition and 0, 20, 40, 60 and 80 minutes after switching to glucose. Whole cell protein extracts were prepared as described previously (Rossmann and Stillman 2013) and analyzed by SDS-PAGE. Western blotting was performed using an anti-Histone H3 antibody (ab1791; Abcam). Ponceau S staining was used to control for variation in protein loading.

### Budding index analysis

Cells were grown in SD –His for 4 hours, treated with α-factor for 2.5 hours, washed once, and released in SD –His at 30°C. Samples were taken every 10 minutes for 1 hour, fixed in 4% paraformaldehyde at room temperature for 15 minutes, washed, and resuspended in KPO_4_/sorbitol buffer (10 mM KPO_4_, 1.2 M sorbitol, pH = 7.5). Brightfield images were captured with a Zeiss AxioImager M2 widefield fluorescence microscope equipped with a 63x Plan-Apochromat (1.4 NA) oil-immersion objective (Zeiss), and a Camera Axiocam 702 mono (Zeiss). The number of budding cells among around 100 imaged cells per strain were counted (**Fig. S7B**).

### dNTP measurement

To measure dNTP levels (**Fig. S7C-D**), ∼25 OD_600_-units of asynchronous cells growing at 26°C or 37°C were harvested on nitrocellulose filters, resuspended immediately in an ice-cold lysis solution containing 12% TCA and 15 mM MgCl_2_, and snap frozen in liquid nitrogen. Cell suspensions were thawed on ice, vortexed briefly, and mixed on an Intelli-Mixer for 15 minutes at 4°C. Cell lysates were then centrifuged at 14000 rpm for 2 minutes at 4°C. The supernatants were neutralized with a dichloromethane-trioctylamine mix and processed for analysis by HPLC as described previously (Jia et al. 2015).

### P-body analysis

To visualize P-bodies (**Fig. 6**), DHH1-yEGFP tagged strains were grown to mid-log phase in YPD. Oxidative stress was induced by addition of 1 M sorbitol for 15 minutes. Cells were fixed in 4% paraformaldehyde at room temperature for 15 minutes, washed, and resuspended in KPO_4_/sorbitol buffer (10 mM KPO_4_, 1.2 M sorbitol, pH = 7.5). Images were acquired on a Zeiss AxioImager M2 widefield fluorescence microscope equipped with ApoTome 2, a 63x Plan-Apochromat (1.4 NA) oil-immersion objective (Zeiss), a Camera Axiocam 702 mono (Zeiss), and an HXP 120 metal-halide lamp used for excitation. 21 focal steps of 0.25 μm were acquired with an exposure time of 1,000 ms using a GFP/YFP 488 filter (excitation filter: 470/40 nm, dichroic mirror: 495 nm, emission filter: 525/50 nm). Images were recorded using ZEN Blue software and analyzed with Fiji.

### ^35^S-methionine incorporation

^35^S-methionine incorporation (**Fig. S8B**) was measured as described previously (Esposito and Kinzy 2014), with the exception that cells were inoculated in 25 mL SD –Met.

## DATA AVAILABILITY

All whole-genome sequencing data are publicly available at NCBI’s Sequence Read Archive (http://www.ncbi.nlm.nih.gov/sra) under accession number PRJNA1131954. All other data are available in the supplementary data files. Data S9 lists all yeast strains and plasmids, the figures in which they were used, and the sources from which they can be obtained. All plasmids generated as part of this study are available from Addgene, all constructed yeast strains are available upon request. UMass Chan Medical School requires completed Material Transfer Agreements for all reagent requests.

## ACKNOWLEDGEMENTS

We thank Sabine van Schie for critical reading of the manuscript and Mika Wesley for help with the Addgene plasmid deposit. This work was supported by grants from the Swiss National Science Foundation (PCEGP3_181242 and 10003570 to J.v.L.), Wellcome (220540/Z/20/A to L.P.), the Swedish Research Council (2022–00675 to A.C.), the Swedish Cancer Society (22 2377 Pj to A.C.), the French National Research Agency (ANR-20-CE13-0010 to G.L.), and the Biotechnology and Biological Sciences Research Council (BB/V015109/1 to M.A.), and a Ramon y Cajal fellowship (RYC-2017–22959 to C.P.)

