## Supplementary Figures for "The modifiers that cause changes in gene essentiality"

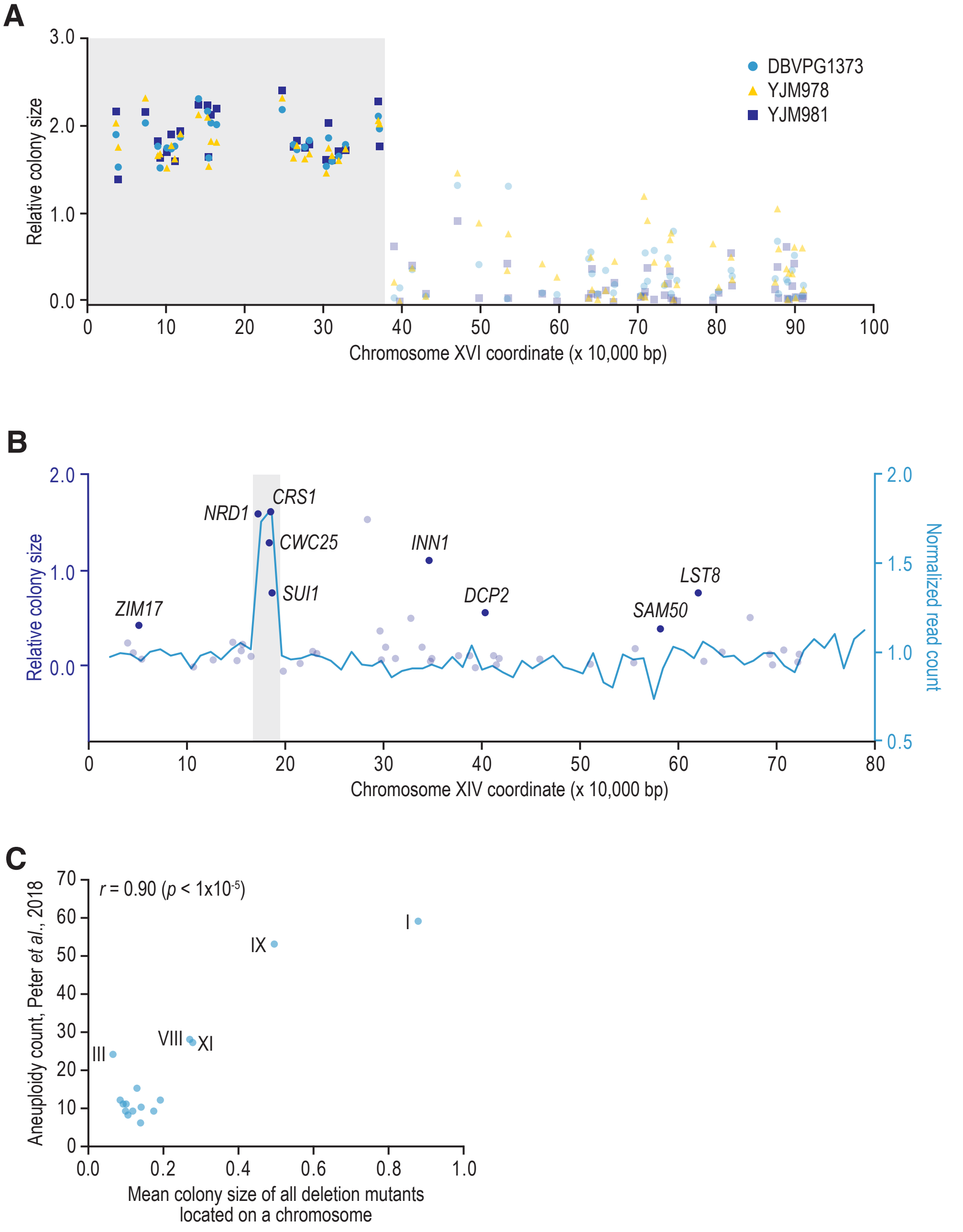


**Fig. S1. Systematic identification of genetic context-dependent essential genes.** (**A**,**B**) Structural variants can suppress essential gene deletion mutants by providing a second (intact) copy of the essential gene. (A) Relative colony size of essential gene deletion mutants located on chromosome XVI in haploid progeny of a cross between S288C and the indicated strains. DBVPG1373, YJM978, and YJM981 carry a translocation between the promoter regions of *ECM34*, located on the left arm of chromosome VIII, and *SSU1*, located on the left arm of chromosome XVI. Twenty-five percent of the haploid progeny of a cross between a strain with the translocation and S288C will have two copies of a large part of the left arm of chromosome XVI (Parts *et al.*, 2021). Essential genes can be deleted on one copy of this arm without loss of viability due to the presence of the second copy. (B) Normalized sequencing read count across 10,000 bp segments of chromosome XIV of CLIB413_1b (cyan) and relative colony size of essential gene deletion mutants in haploid progeny of a cross between S288C and CLIB413_1b (blue). CLIB413_1b carries a duplication on the left arm of chromosome XIV that, similar to the translocation described in (A), permits deletion of essential genes in the affected region without loss of viability by providing a second copy of the genes. Sequencing read counts at the telomeres are not shown due to a high variability in counts in these regions. Genes that are essential in S288C but were nonessential in the cross progeny are highlighted in darker colors. Grey shading indicates the duplicated regions. (**C**) Correlation between aneuploidy prevalence and viability of essential gene deletion mutants. The frequency of aneuploidies for each of the 16 yeast chromosomes in a set of 1,011 natural yeast isolates (Peter *et al.*, 2018) is plotted against the mean colony size for all essential gene deletion mutants located on that chromosome in the wild strain crosses. Pearson's correlation coefficient and the corresponding *p* value are indicated.

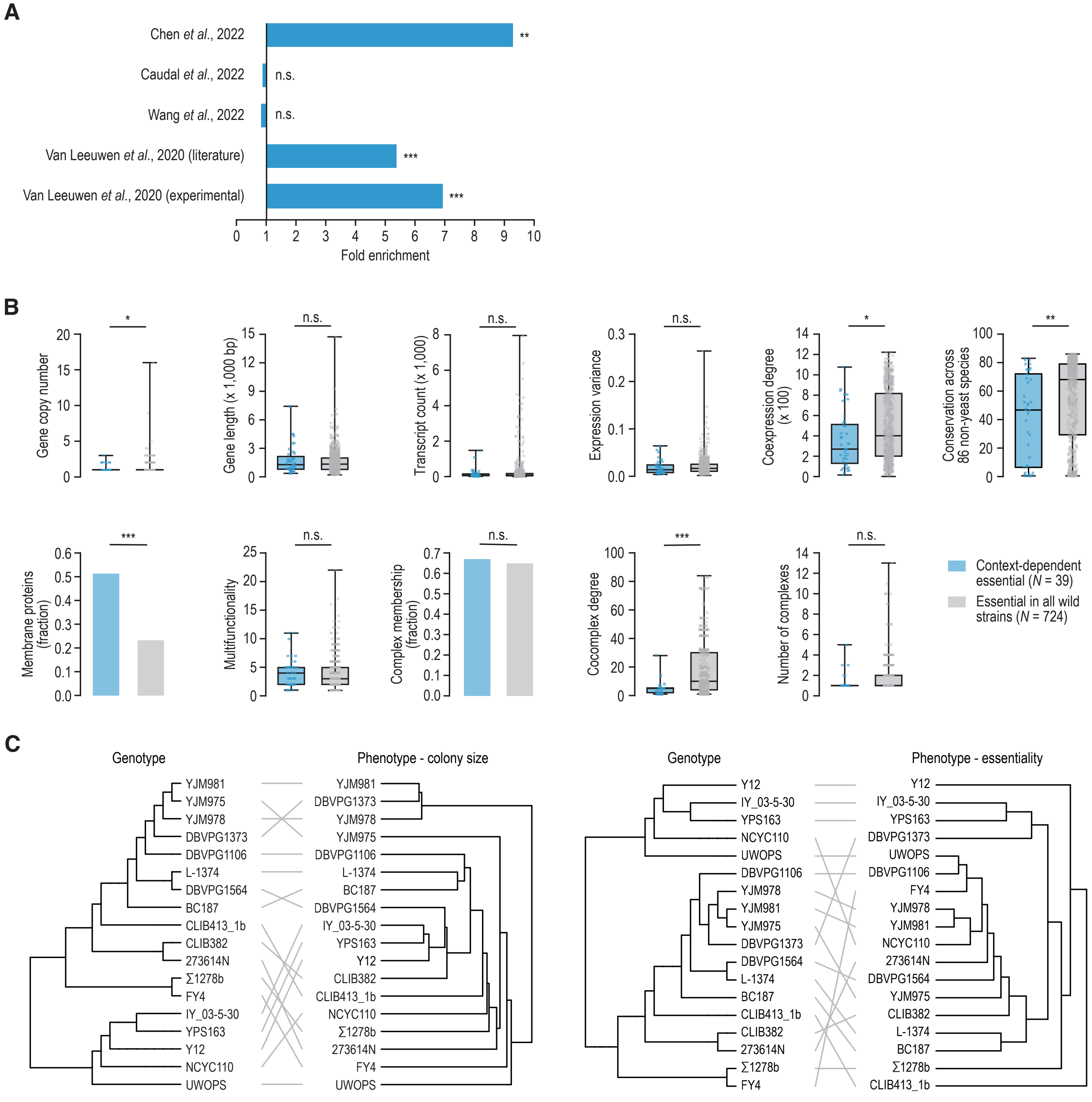


**Fig. S2. Properties of genetic context-dependent essential genes.** (**A**) Fold enrichment of context-dependent essential genes identified in our study, for similar gene sets identified in other studies: "Polymorphic genes" (Chen *et al.*, 2022), "Background healthy genes" (Caudal *et al.*, 2022), "Strain-specific genes" (Wang *et al.*, 2022), and "Dispensable essential genes" (Van Leeuwen *et al.*, 2020). No significant overlap was seen between our study and studies looking at the effect of gene inactivation on fitness, rather than essentiality, across yeast strains (Caudal *et al.*, 2022; Wang *et al.*, 2022). (**B**) Comparison between context-dependent essential genes and genes that were essential in all genetic backgrounds for various gene- and protein-level properties. Observed enrichments of context-dependent essential genes are comparable to those of "Dispensable essential genes" described previously (Van Leeuwen *et al.*, 2020; Pons and Van Leeuwen, 2023). (**C**) Genotype and phenotype trees are somewhat concordant. Genotype: clustering of the wild strains used in this study based on genomic variants. Phenotype: hierachical clustering of either the normalized colony sizes of haploid segregant populations lacking the essential gene (left; Data S2) or of gene essentiality profiles (right; Data S4). IY_03-5-30 = IY_03-5-30-1-1-1_(1); UWOPS = UWOPS87-2421. Statistical significance was determined using Fisher's exact or Mann-Whitney U tests. * $p < 0.05$, ** $p < 0.005$, *** $p < 0.0005$, n.s. = not significant.

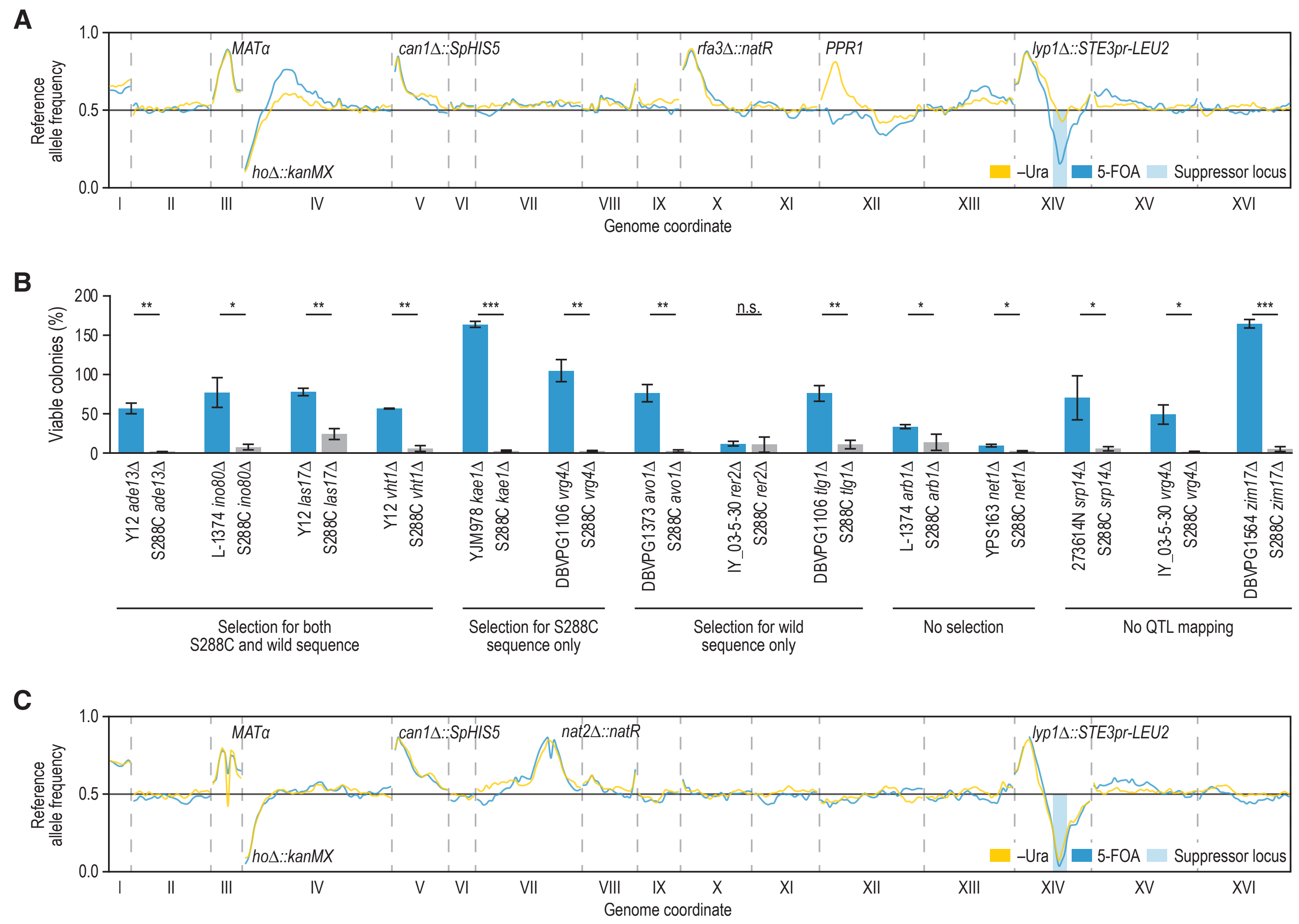


**Fig. S3. Mapping suppressor loci by sequencing segregant pools.** (**A**) The *PPR1* locus of S288C is selected in –Ura samples of crosses with CLIB413_1b. Reference allele frequency along the yeast genome in *rfa3*Δ progeny of a cross of an *rfa3*Δ [*RFA3*, *URA3*] S288C strain to CLIB413_1b, either in the presence of the plasmid carrying *RFA3* (–Ura, yellow) or in the absence of the plasmid (5-FOA, cyan). Loci carrying selection markers that were used during the mapping procedure are labeled. The S288C strain that was used in the cross carried an aneuploidy of chromosome I (Data S11), which is visible as an increase in reference allele frequency for this chromosome. (**B**) Bypass suppression is generally driven by natural variants present in the wild genomes. Indicated strains were transformed with a plasmid encoding Cas9 as well as a guide RNA targeting the essential gene, and a repair template that when integrated into the genome introduced a frameshift in the targeted gene. Plotted are the mean number of viable colonies obtained for each essential gene in both the wild background and the reference strain S288C, as percentage of the number of colonies obtained when a nonessential gene (*YFR054C*) was targeted. Genes and wild strains are sorted by their phenotype in the bulk segregant analysis (Fig. 3): crosses that showed regions of selection for both S288C and wild sequence; crosses that showed regions of selection for either S288C or wild sequence only; crosses that showed no regions with significant selection for either wild or S288C sequence; and crosses for which we did not perform bulk segregant analysis. Error bars represent the standard error of the mean of two independent experiments. * $p < 0.05$, ** $p < 0.005$, *** $p < 0.0005$, n.s. = not significant, unpaired one-tailed t-test. IY_03-5-30 = IY_03-5-30-1-1-1_(1). (**C**) Reference allele frequency as described in (A), but for *nat2*Δ progeny of a cross between a *nat2*Δ [*NAT2*, *URA3*] S288C mutant and IY_03-5-30-1-1-1_(1). The wild allele of *MKT1*, which is located in the middle of the highlighted suppressor locus on chromosome XIV, is strongly selected in both the –Ura and 5-FOA samples. The S288C strain that was used in the cross carried an aneuploidy of chromosome I (Data S11), which is visible as an increase in reference allele frequency for this chromosome.

**A**

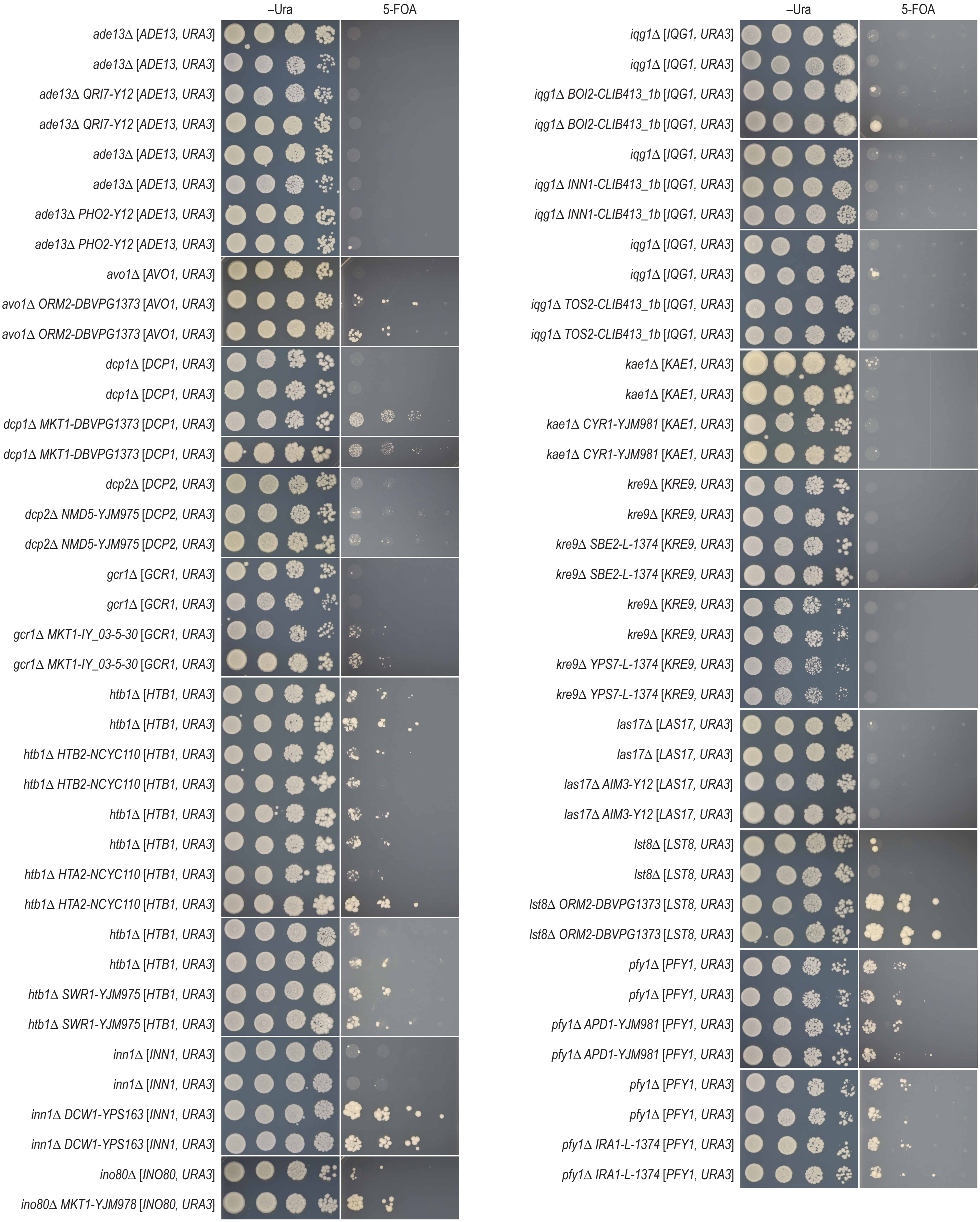


Continued on the next page

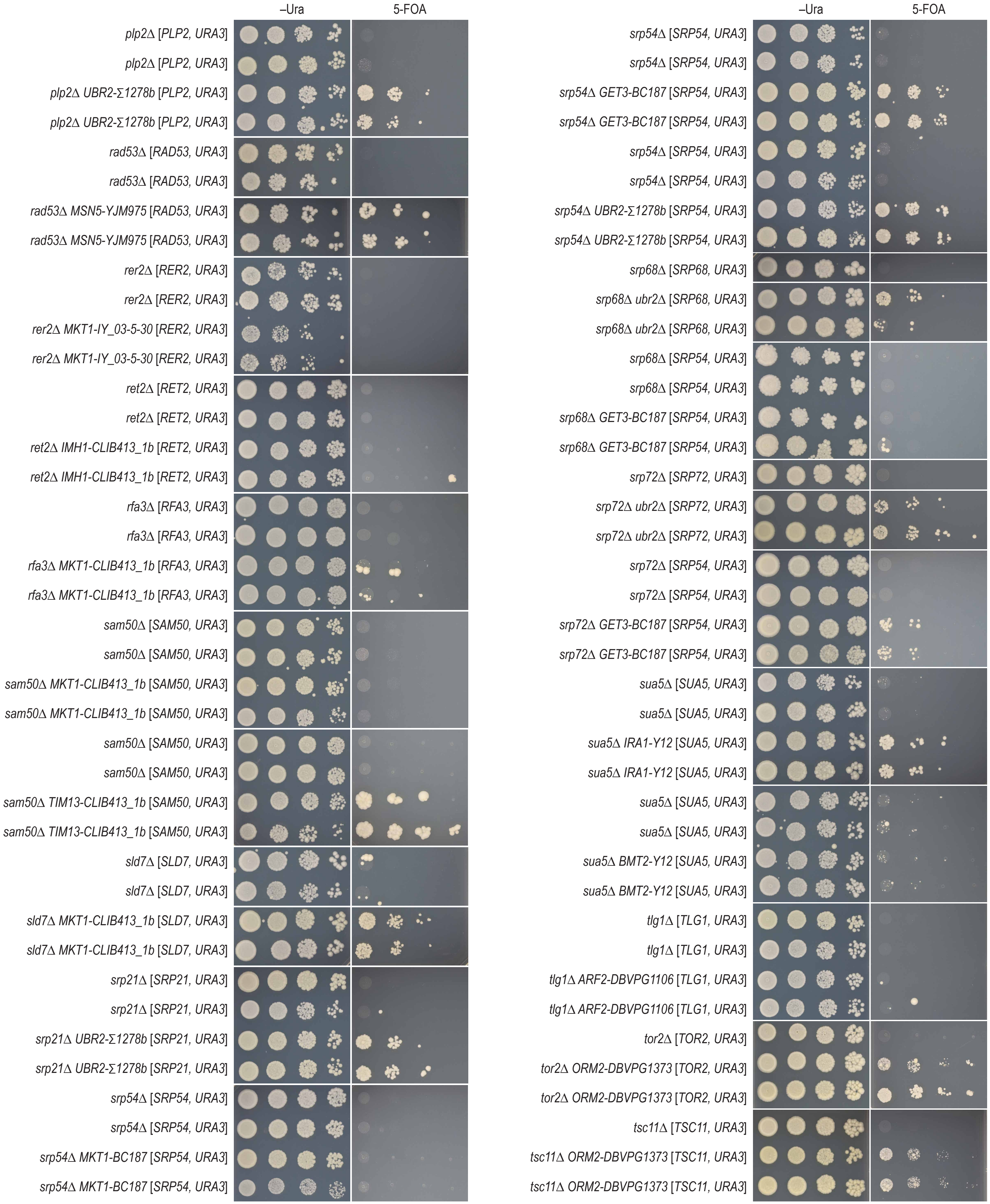


Continued on the next page

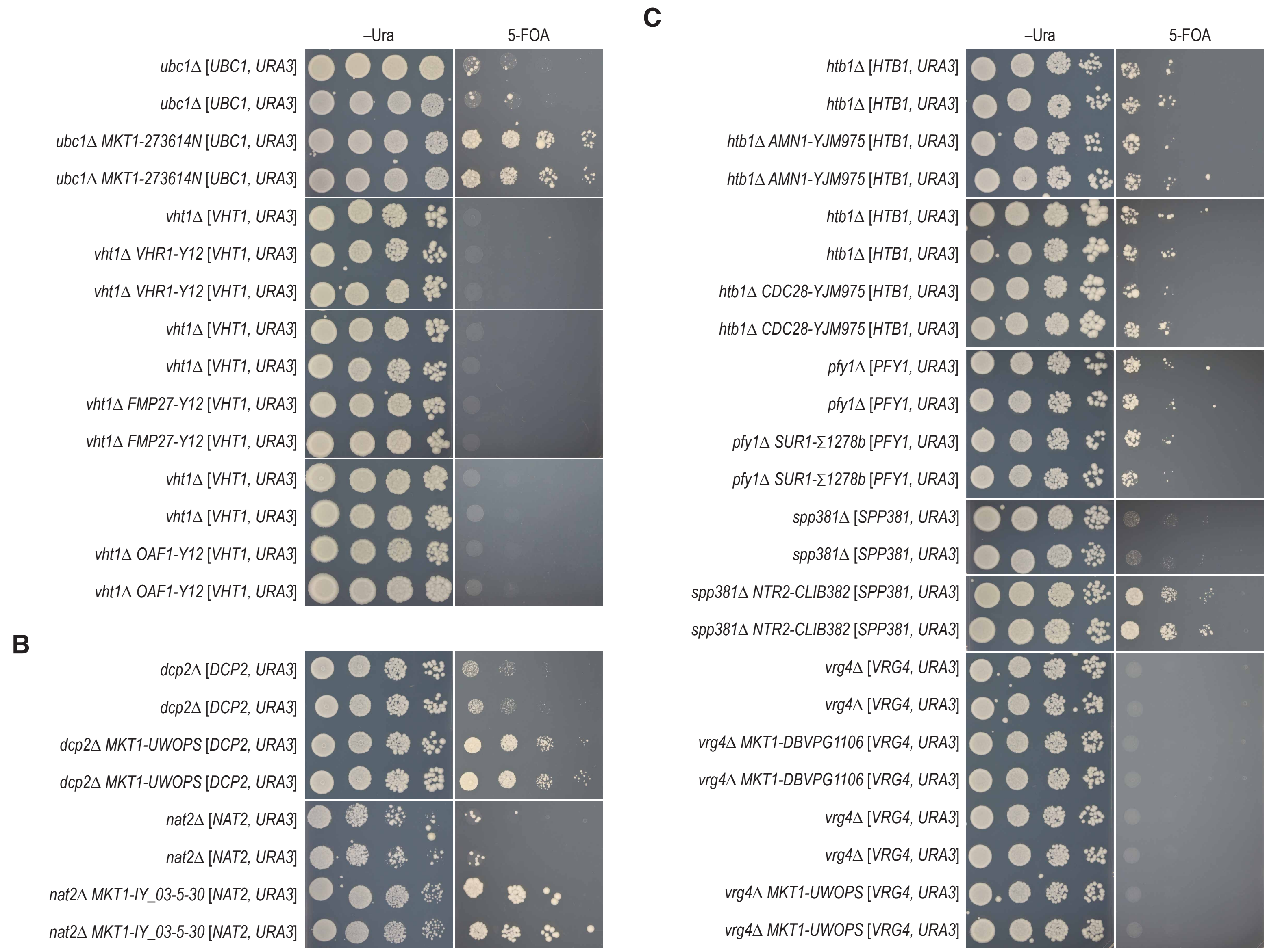


**Fig. S4. Suppressor gene validation.** (**A-C**) To validate the suppressor candidates, we replaced the reference alleles of the potential suppressor genes with their wild versions in the reference strain background, and tested for suppression of the corresponding essential gene deletion mutant. Cultures of the indicated strains were grown until saturation, and a series of five- or ten-fold dilutions was spotted on agar plates, either selecting for the plasmid carrying the essential gene (–Ura) or selecting against the plasmid (5-FOA), and incubated at 26°C for six days. (A) Allele swaps for genes that were located in mapped QTLs. For *rad53Δ MSN5*, the *MSN5* S288C and YJM975 alleles were expressed from plasmid and *MSN5* was deleted in the genome. For *srp68Δ UBR2* and *srp72Δ UBR2*, we used an *UBR2* deletion mutant as the *UBR2-Σ1278b* allele carries a premature stop codon and is thus likely a loss-of-function allele. Moreover, for *ino80Δ MKT1*, *pfy1Δ IRA1*, *srp68Δ GET3*, and *srp72Δ GET3*, the alleles that were used in the allele swaps came from a different wild strain than that was used in the QTL analysis, but that carried similar nonsynonymous variants in the swapped gene. Full genotypes of all used strains can be found in Data S9. (B) Additional allele swaps for genes that were located within the query linkage group (*DCP2*) or that were strongly selected both in the presence and absence of the essential gene plasmid (*NAT2*). (C) Allele swaps for genes located in loci showing weak selection for wild strain sequence in the 5-FOA sample, that did not meet the selection criteria used for QTL calling. IY_03-5-30 = IY_03-5-30-1-1-1_(1); UWOPS = UWOPS87-2421.

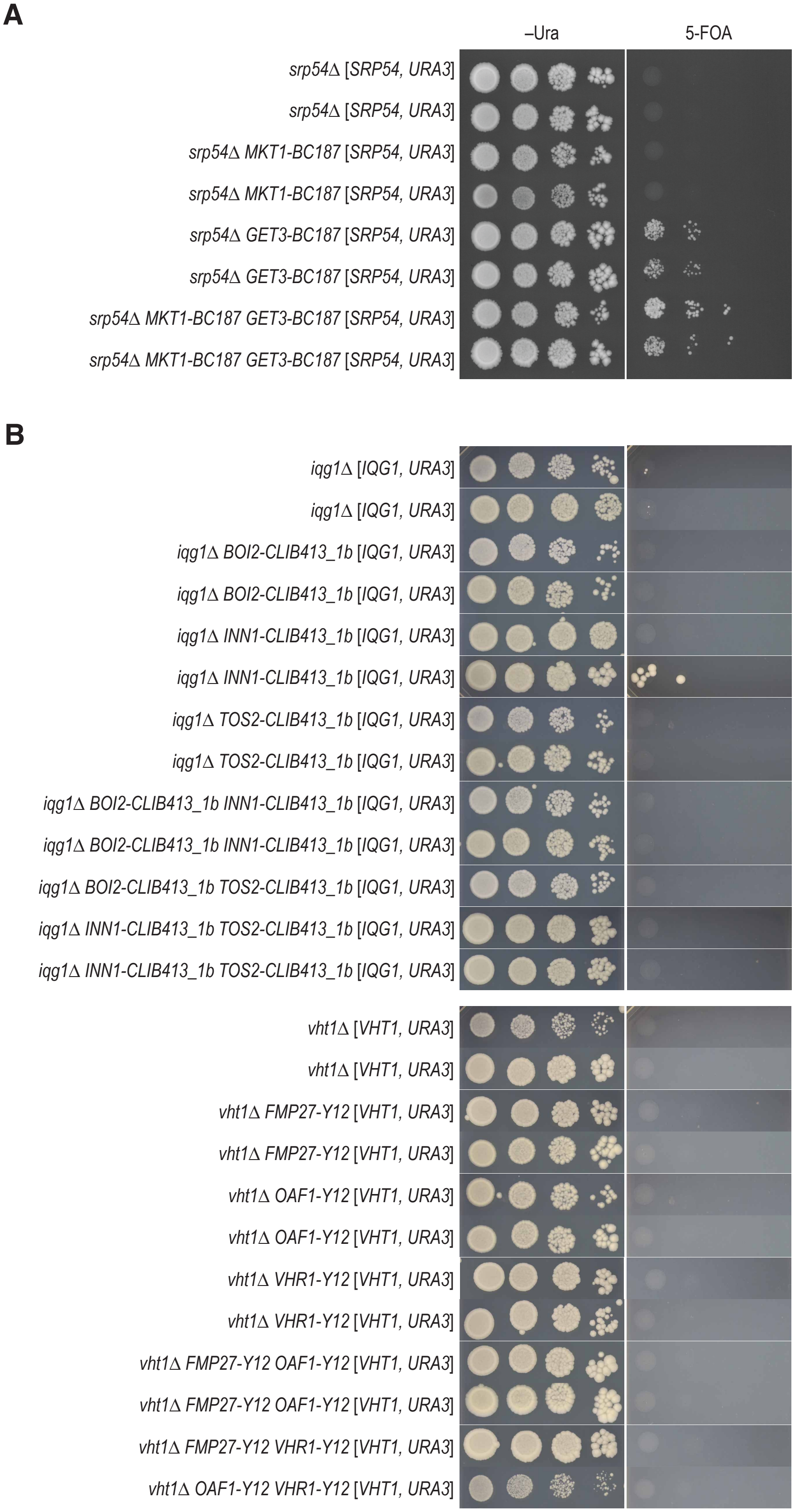


**Fig. S5. Bypass suppressors act in isolation.** (**A**,**B**) To test whether multiple genes contributed to the bypass suppression, we generated strains in which the reference alleles of two suppressor candidate genes were replaced with their wild versions in the reference strain background, and tested for suppression of the corresponding essential gene deletion mutant. Cultures of the indicated strains were grown until saturation, and a series of ten-fold dilutions was spotted on agar plates either selecting for the plasmid carrying the essential gene (–Ura) or selecting against the plasmid (5-FOA) and incubated at 26°C for six days. (A) Double allele swaps for *srp54*Δ candidate suppressor genes. The BC187 allele of *GET3* could bypass *srp54*Δ, but the *MKT1-BC187* allele could not (see also Fig. S4). Combination of both wild alleles slightly improved the fitness compared to *srp54*Δ *GET3-BC187*. (B) Double allele swaps for *iqg1*Δ and *vht1*Δ candidate suppressor genes. None of the candidate suppressors could bypass the essential gene deletion mutant either individually (see also Fig. S4) or in pairwise combinations.

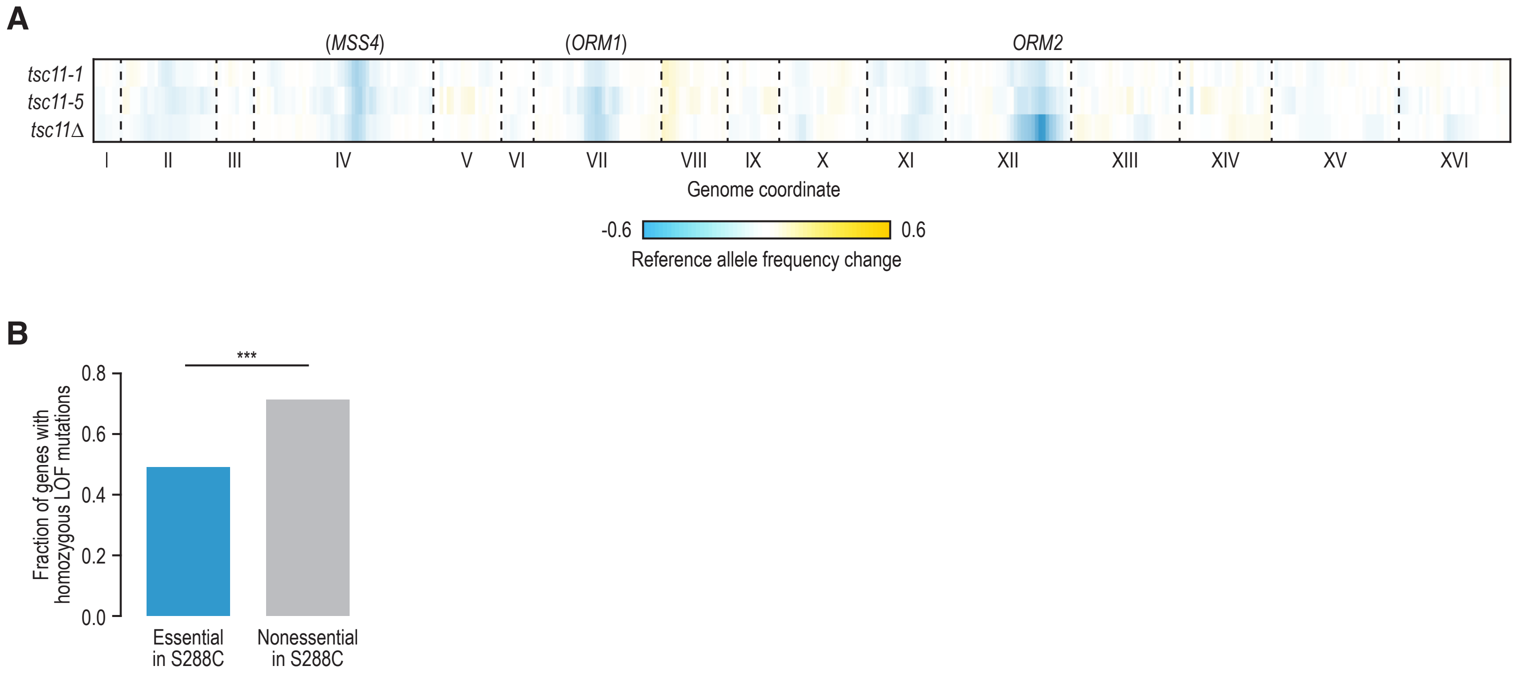


**Fig. S6. Characterization of natural bypass suppression variants.** (**A**) Mapping results for segregant pools of crosses between the indicated *TSC11* alleles and DBVPG1373. The change in S288C allele frequency between meiotic progeny isolated in the presence and the absence of the plasmid carrying *TSC11* (for *tsc11*Δ) or between 26°C and 34°C (for *tsc11-1* and *tsc11-5*; Parts *et al*., 2021) is plotted by genomic coordinate. Causal suppressor genes are indicated for regions that show selection for DBVPG1373 sequence. Genes in brackets have not been validated experimentally. (**B**) Fraction of all yeast genes that are either essential (N = 1,071) or nonessential (N = 4,800) in S288C that carried homozygous loss-of-function (LOF) mutations in one or more of 1,011 natural yeast isolates (Peter *et al*., 2018).

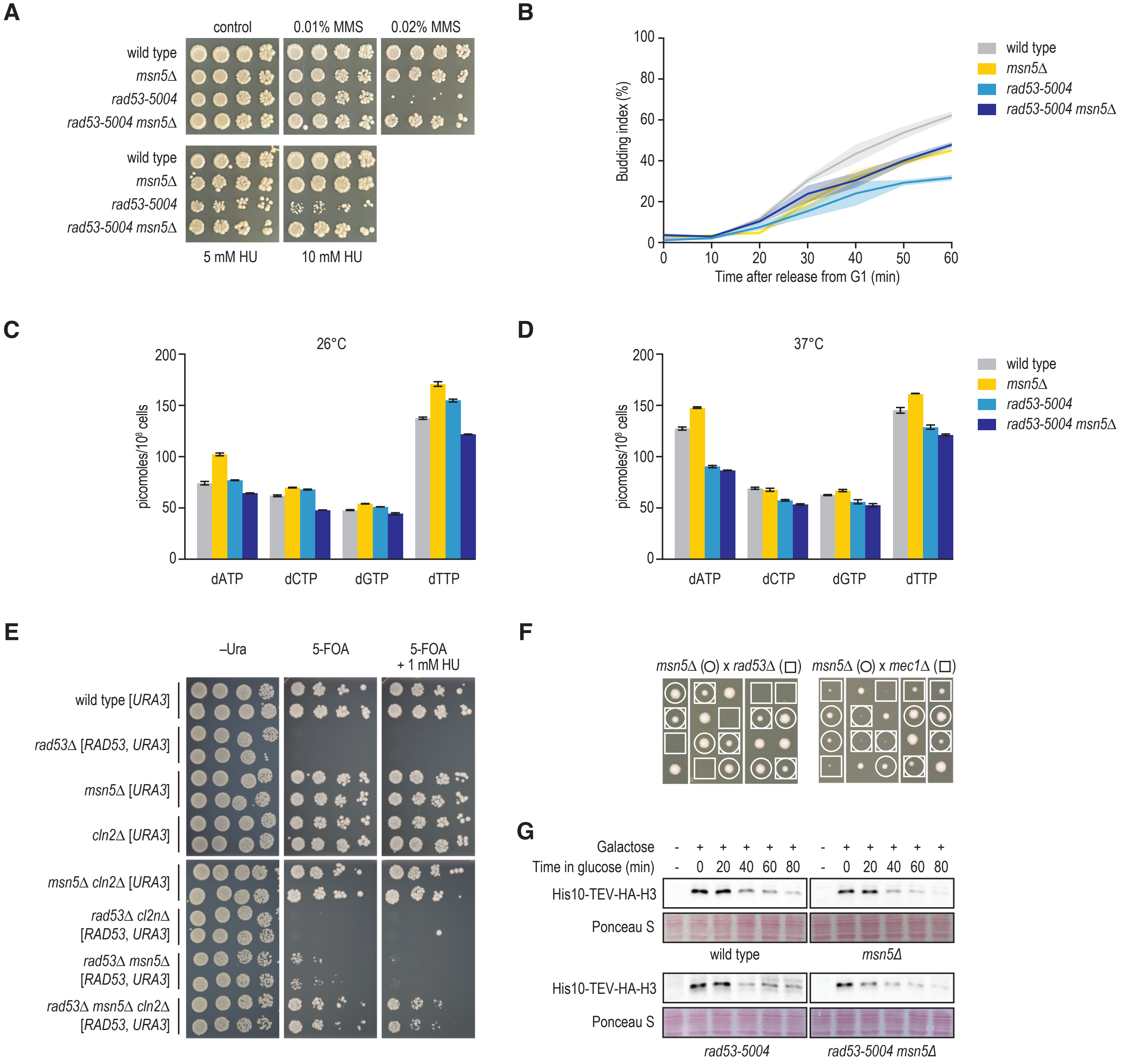


**Fig. S7. Loss-of-function mutations in *MSN5* can suppress the lethality of a *rad53*Δ mutant.** (**A**) The *rad53-5004* allele is a partial loss-of-function allele that leads to increased MMS and HU sensitivity, which can be suppressed by *msn5*Δ. Cultures of the indicated strains were diluted to an optical density at 600 nm of 1, and a series of 10-fold dilutions was spotted on agar plates and incubated at 30°C for three days. (**B**) Deletion of *MSN5* does not slow down cell cycle progression of *rad53-5004* cells. Cultures of the indicated strains were synchronized with mating pheromone, released from G1, and the percentage of budding cells was monitored at 30°C. Although cell cycle phase progression was delayed in both *rad53-5004* and *msn5*Δ strains compared to the wild type, deletion of *MSN5* in *rad53-5004* rescued cell cycle progression up to the levels of the *msn5*Δ single mutant. Shading represents the standard deviation of three independent measurements. (**C**-**D**) Absence of *MSN5* does not increase dNTP levels in *rad53-5004* strains. dNTP levels were measured in duplicate in asynchronous cells of the indicated strains at 26°C (C) or 37°C (D). Error bars represent the standard error of the mean. (**E**) Suppression by *msn5*Δ does not involve *CLN2*. Cultures of the indicated strains were diluted to an optical density at 600 nm of 1, and a series of 10-fold dilutions was spotted on agar plates and incubated at 30°C for three days. (**F**) Deletion of *MSN5* does not suppress the severe fitness defect of *mec1*Δ mutants. Strains deleted for *MSN5* were crossed to strains deleted for either *MEC1* or *RAD53* (covered with a plasmid). The resulting diploids were cured of the plasmid, strains were sporulated, and tetrads were dissected. Images were taken after three days of incubation at 30°C. Part of this figure is also shown in Fig. 5A. (**G**) Histone degradation is restored in *rad53-5004 msn5*Δ double mutants. Wild-type, *msn5*Δ, *rad53-5004*, and *rad53-5004 msn5*Δ strains expressing a galactose inducible, His10-TEV-HA-tagged histone H3 gene (Gunjan and Verreault, 2003) were cultured in media containing galactose to overexpress histone H3. Cells were then switched to media containing glucose and harvested every 20 min. The levels of tagged H3 were determined by Western blotting. The cells were blocked and kept in G1 using α-factor for the duration of the experiment.

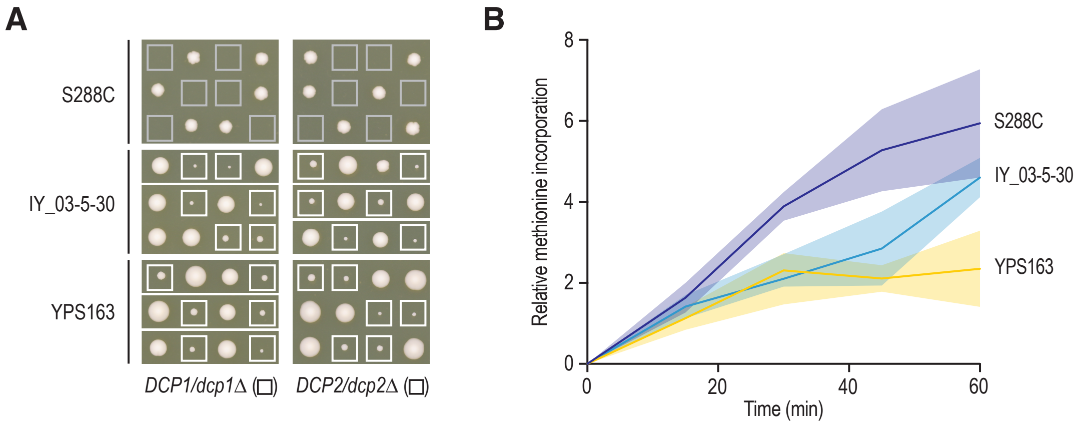


**Fig. S8. The Dcp1-Dcp2 decapping enzyme complex is not required for viability in most natural yeast isolates.** (**A**) Diploid strains with the indicated genetic background and carrying a heterozygous deletion allele of either *DCP1* or *DCP2* were sporulated and tetrads were dissected. (**B**) Translation rates of the indicated strains were measured as the relative incorporation of [$^{35}$S]methionine into proteins over time. Plotted are the means of four independent experiments, shading indicates the standard deviation. IY_03-5-30 = IY_03-5-30-1-1-1_(1).
